# Sensory Adaptations: Insights into the Vomeronasal System of the Iberian Wolf

**DOI:** 10.1101/2023.09.14.557816

**Authors:** Irene Ortiz-Leal, Mateo V. Torres, José-Daniel Barreiro-Vázquez, Ana López-Beceiro, Luis Fidalgo, Pablo Sanchez-Quinteiro

## Abstract

Wolves, like other canids, extensively use chemical signals for various aspects of communication, including territory maintenance, reproductive synchronization, and social hierarchy signaling. Pheromone-mediated chemical communication operates unconsciously among individuals, acting as a mysterious sixth sense that regulates both their physiology and behavior. Despite their crucial role in the life of the wolf, there is a surprising lack of comprehensive research on the neuroanatomical and physiological bases of chemical communication in wolves.

This study delves into the Iberian wolf vomeronasal system (VNS) and examines potential changes brought about by dog domestication. Our findings show that the Iberian wolf possesses a fully functional VNS vital for pheromone-mediated communication. While macroscopic similarities between the wolf and domestic dog VNS are observed, there are notable microscopic differences. These include the presence of neuronal clusters associated with the sensory epithelium of the vomeronasal organ (VNO) and a higher differentiation degree of the accessory olfactory bulb (AOB). Immunohistochemical markers reveal the expression of the two main families of vomeronasal receptors (V1R and V2R) in the VNO. However, only the V1R family is expressed in the AOB.

These findings not only provide deep insights into the VNS of the wolf but also hint at how domestication might have altered neural configurations that underpin species-specific behaviors. This understanding has implications for innovative strategies, such as employing semiochemicals for wolf population management, aligning with modern conservation goals.

## INTRODUCTION

After generations of a negative interaction between man and wolf that resulted in the demise of wolf populations in the majority of Europe, new social attitudes toward wildlife and conservation have emerged, institutionalizing wolf protection (Stohr and Coimbra 2013). Nevertheless, wolf population control remains a difficult and controversial endeavor. As wolves spread into agricultural regions, human-wolf conflict intensifies. Due to the wolf great reproductive capacity and tendency to wander there are few areas where wolves might be reintroduced without some type of control (Mech 1995). The use of deadly measures to manage wolves is becoming less acceptable, but they are still the most effective way to reduce the harm that wolves cause to cattle and pets, at least until non-lethal alternatives become available (Wielgus and Peebles 2014). Aversive conditioning (Gustavson and Nicolaus 1987) has not yet been demonstrated to be successful with wild wolves (Fritts et al. 1992). However, the employment of semiochemicals, either pheromones or kairomones (Fortes-Marco et al. 2015), as a method for population control is a potential tool that has received only a limited amount of investigation (Petrulis 2013; Van Den Berghe et al. 2019; Riddell et al. 2021). Chemical signals play a significant role in intraspecific communication, acting as a mediator between the sexual behavior of the species and the physiological processes involved in reproduction. In the case of the wolf, the species deploys scent marks for the purposes of maintaining its territory (Peters and Mech 1975; Barja et al. 2004), establishing reproductive synchrony (Rothman and Mech 1979), and expressing its social position (Barja et al. 2008). Within a group, scent marking is restricted to mating males and females, and subordinates do not display it unless they are vying for dominance (Asa et al. 1990). Social suppression of reproduction is typical of wild canids (Macdonald et al. 2019), but it also occurs in rodents like the naked mole-rat, where it is mediated by semiochemicals sensed via a functional vomeronasal organ (VNO) (Dennis et al. 2020).

As a preeminent symbol among endangered species, wolves may be one of the most studied wild mammals in the world. However, beyond of the study of territory-related pheromones in urine and faecal marks (Barja et al. 2004; Barja et al. 2008; Raymer et al. 1985; Raymer et al. 1986; Wirobski et al. 2021), there is a surprising lack of information on the neuroanatomical and physiological basis of wolves chemical communication, for both the olfactory and vomeronasal systems, parts of the sensory system responsible for the processing of semiochemicals. To the best of our knowledge, there is no description of the vomeronasal system (VNS) of the wolf, and the information that is available on the main olfactory system (MOS) is restricted to the comparative morphological and neurochemical research of the main olfactory bulb (MOB) of domestic and wild canids (Ortiz-Leal et al. 2022a).

The MOS is made up of millions of neuroreceptor cells that are placed in the olfactory epithelium (OE) that covers the olfactory concha (Barrios et al., 2014; Bressel et al., 2016). These cells are responsible for transmitting information to the MOB (Doucette et al. 1983; Su et al. 2009). Because of its close connection to the limbic system, the MOS has been linked to both memory and the awareness of conscious sensations (Slotnick 2001; Ubeda-Bañon et al. 2011). In contrast, the VNO, sensorial component of the VNS (Kratzing 1971; Tomiyasu et al. 2022), comprises neuroreceptor cells that send their signals through the vomeronasal nerves (NVN) (McCotter 1912; Smith et al. 2015) to the accessory olfactory bulb (AOB) (Frahm and Bhatnagar 1980; Mohrhardt et al. 2018). The VNS is specialized for detecting pheromones (Powers and Winans 1975; Kunkhyen et al. 2017), kairomones (Isogai et al. 2011; Fortes-Marco et al. 2013), and molecules of the major histocompatibility complex (Leinders-Zufall et al. 2000; Leinders-Zufall et al. 2014). Among the numerous functions of the VNS are playing non-conscious roles in socio-sexual behaviours (Baum and Cherry 2015; Abellán-Álvaro et al. 2022), maternal recognition (Kohl et al. 2017; Navarro-Moreno et al. 2020), sickness avoidance behaviour (Boillat et al. 2015; Bufe et al. 2019) and the detection of predators (Tsunoda et al. 2018).

Despite the fact that extrapolations are known to be problematic in the study of the neuroanatomy of the VNS (Salazar and Sánchez-Quinteiro 2009; Salazar et al. 2016), the Rodentia Order has served as the major referent for the VNO research carried out on mammalian species (Salazar et al., 2013). Thus, the neuroanatomical data available on the VNS of canids is mainly focused in the domestic dog VNO (Dennis et al. 2003; Salazar et al. 2013; Mahdy et al. 2019) and AOB (Jawlowski 1956; Miodonski 1968; Salazar et al. 1994b; Nakajima et al. 1998), as well as its secondary projections to the vomeronasal amygdala (Miodonski 1968). As far as wild canids are concerned, until recently the majority of available information was restricted to the part devoted to the rhinencephalon in the study of the brain of the African wild dog *Lycaon pictus* by Chengetanay et al. (2020) and the studies on the fox vomeronasal amygdala (Równiak and Bogus-Nowakowska 2020; Równiak et al. 2022). However, the recent thorough investigation of the VNO, MOB, AOB, and olfactory limbus of the red fox (Ortiz-Leal et al. 2020; Ortiz-Leal et al. 2022a; Ortiz-Leal et al. 2022b) revealed that this species possesses a neuroanatomical structure that is significantly dissimilar to the pattern reported in the canine VNS. In dogs, the restricted development of the VNO epithelium and the inadequate differentiation of the glomerular and nerve layers of the AOB, with a small size and the absence of the characteristic cytoarchitecture found in other mammalian species (Meisami and Bhatnagar 1998) are notorious. In contrast, fox investigations reveal a VNS with a high degree of differentiation and the existence of features unreported in the dog, particularly at the VNS level. Likewise, in the case of the African wild dog, the degree of development shown by the AOB points to an enhanced sensitivity compared to the domestic dog (Chengetanai et al. 2020). All these anatomical differences support the current hypothesis that the domestication of the dog has resulted in an involution of olfactory development associated with the detection of pheromones and other semiochemicals by the VNS.

It is however, still an open question whether throughout the approximately 10,000 years that phylogenetically separate the wolf and the dog (Graphodatsky et al. 2008; Bergström et al. 2022), the intense selection pressure associated with domestication may have led to the appearance of modifications in the configuration of neural structures that support species-specific behaviors such is the case of the VNS. Therefore, this study aims to not only address the important gap in the morphological and immunohistochemical knowledge of the vomeronasal system of the wolf, but also to shed light on how the domestication process may have influenced the organization of the central nervous system. We have used various tissue dissection and microdissection techniques, and computed tomography (CT) imaging, followed by general and specific histological stainings, including immunohistochemical and lectin-histochemical labelling techniques.

Among the variety of antibodies used for the immunohistochemical study of the wolf VNS, special attention should be paid to the study of the pattern of expression of G-proteins in the sensory epithelium of the VNO, the vomeronasal nerves, and the AOB. The immunohistochemical characterization of both Gαi2 and Gαo G-proteins has been considered to serve as an excellent phenotypic indicator of the expression in the VNS of the two main families of vomeronasal receptors, V1R and V2R respectively. The effective expression of both G-proteins is a feature observed in Rodents, such as mouse, rat, octodon (Suárez and Mpodozis 2009), guinea pig (Takigami 2004), and capybara (Suárez et al. 2011b; Torres et al. 2020); Lagomorpha, such as rabbit (Villamayor et al. 2020); Marsupials, including opossum (Halpern et al. 1995) and wallaby (Torres et al. 2022); and tenrecs (Suárez et al. 2009). However, in other mammals, such as the dog, cat, sheep and goat, the differential expression of G proteins and vomeronasal receptors has not been observed, as these species exclusively express the V1R receptor family (Takigami 2004; Salazar et al. 2007; Salazar et al. 2013; Salazar and Sánchez-Quinteiro 2011). The absence of V2R receptors has been theorised to be the result of the domestication process, during which artificial selection may have produced an involution of the VNS in canids (Barrios et al., 2014; Jezierski et al., 2016). Therefore, investigating the expression patterns of these receptors and the general anatomy of the VNS in a wild canid with close phylogenetic proximity to the dog, such as the wolf, could contribute to a deeper comprehension of this theory.

## METHODS

In this study, we utilized a sample of five adult male wolves. These wolves originated from wildlife recovery centers in the provinces of Galicia and were unfortunately involved in fatal traumatic accidents. Only those that had died recently and displayed no external or internal head injuries were included in our research. All samples were used with the compulsory permissions by the Galician Environment, Territory and Tenement Council (CMATV approval numbers EB-009/2020 and EB-007/2021).

All the heads were dissected as soon as they arrived to the Faculty of Veterinary, unless one head that was frozen and transverse sectioned, to compose a macroscopic photographic series. The rest of the heads were dissected extracting the whole brains after opening dorsally the cranium and removing the lateral walls of both the cranial cavity and the ethmoidal fossa with the help of an electric plaster cutter and a gouge clamp. The VNOs were identified after removing the nasal bones and the lateral walls of the nasal cavity. The bone tissue surrounding the VNO ventrally and medially was dissected from all samples, unless one sample which was decalcified for two weeks week to microscopically study the topographic relationship of the VNO with the incisive duct. The decalcifying agent used was Shandon TBD-1 Decalcifier (Thermo, Pittsburgh, PA), and it was applied while stirring continuously at room temperature.

All the samples were fixed in Bouin’s fluid for 24 h., then transferred to 70% ethanol, embedded in paraffin, and cut by a microtome. The olfactory bulbs were cut transversely and sagittaly by a microtome with a thickness of 8 µm, whereas the VNO samples were sectioned with a thickness of 6-7 µm. The VNO was serially transverse sectioned along its entire length, from caudal to rostral. The slides were stained using haematoxylin-eosin, Periodic acid-Schiff (PAS), Alcian Blue, and Gallego’s Trichome stains (GTr) (Ortiz-Hidalgo 2011). The staining protocols are explained in detail in Salazar et al. (2003) and Torres et al. (2023a).

### Computed Tomography Scans

CT of the head was performed in a 16-slice helical multidetector scanner (Hitachi Eclos 16), obtaining both bone and soft tissue algorithm series in sternal recumbency. For the bone series a 1,25 mm slice thickness every 0,625 mm, while for the soft tissue series a 2,5 mm slice thickness every 1,25 mm was applied. Exposure factors were 120 kVp and 200 mA, with 1 second per rotation and a pitch of 0,5.

### Lectin histochemistry protocol

LEA and BSI-B_4_ were employed as biotinylated conjugates. Deparaffinized and rehydrated slides were incubated in a solution of 3% hydrogen peroxide to quench endogenous peroxidase activity. Following this, the sections were incubated in a 2% solution of bovine serum albumin (BSA) in 0.1 M phosphate buffer (PB) at pH 7.2 for 30 min. Overnight incubation was performed with LEA and BSI-B_4_ lectins, separately, in a 0.5% BSA mixture. After two brief washes in PB, the slides were incubated in avidin-biotin-peroxidase (ABC) complex (Vector Laboratories, Burlingame, CA, USA) at room temperature for 90 minutes. A 0.003% hydrogen peroxide and 0.05% 3,3-diaminobenzidine (DAB) solution in a 0.2 M Tris-HCl buffer were used to visualize the ensuing reaction, which resulted in a brown-colored deposit.

For UEA lectin, the initial two steps mirrored those used for LEA and BSI-B_4_. Slides were subsequently incubated for 60 minutes in a 0.5% BSA–UEA mixture. Then, anti-UEA peroxidase-conjugated antibody was added, and overnight incubation ensued. The reaction was revealed through the application of a DAB solution, as described for the LEA and BSI-B_4_ procedure.

As controls, tests without lectin addition and also pre-absorbed lectins with excessive corresponding sugars, were performed.

### Immunohistochemistry methodology

The first step involved treating all samples with a 3% hydrogen peroxide solution to inhibit endogenous peroxidase. Subsequently, samples were immersed in either a 2.5% horse serum solution, compatible with the ImmPRESS Anti-Mouse/Anti-Rabbit IgG reagent kit (Vector Laboratories, Burlingame, CA, USA), or 2% BSA for half an hour to preclude non-specific binding. Samples were incubated overnight at 4°C with the primary antibodies. The next day, depending on the blocking agent used, samples were incubated for 30 min with either the ImmPRESS VR Polymer HRP Anti-Rabbit IgG or Anti-mouse IgG reagents, with the exception of samples treated with anti-OMP antibodies derived from goats, which were first incubated in a biotinylated anti-goat IgG and afterwards incubated in ABC reagent for 1.5 hours under humid conditions. Three sequential 5-minute PB washes were carried out between steps. Prior to the visualization stage, all samples were rinsed for 10 minutes in 0.2 M Tris-HCl buffer at pH 7.61. DAB chromogen was used for visualizing, following the same protocol used for lectin histochemical labelling. Negative controls omitted the primary antibodies.

### Image Capture and Digital Alterations

Images were digitally captured using a Karl Zeiss Axiocam MRc5 camera coupled with a Zeiss Axiophot microscope. Adobe Photoshop CS4 was employed for brightness, contrast and balance adjustment; however, no enhancements, additions, or relocations of the image features were made. Additionally, an image-stitching software (PTGuiPro) was used for low magnification images composed of several photographs.

## RESULTS

The VNS will be studied at both the macroscopic and microscopic levels. For each of these levels, detailed descriptions of the vomeronasal organ, vomeronasal nerves, and accessory olfactory bulb are provided.

### 1. Macroscopic study of the VNS

#### 1.1 Vomeronasal organ (Fig. 1-5)

As a preliminary step to the dissection of the vomeronasal organ, a cross-sectional macroscopic anatomic study of the nasal cavity was performed on one specimen (Fig. 1A-E). The eight sections chosen extended from the area of the nasal vestibule to the area of the ethmoidal turbinates (Fig. 1C). Among them, the one corresponding to level 6 contained the central part of the VNO (Fig. 1B,D). At higher magnification, the VNO corresponds to two tubular structures located in the ventral part of the nasal septum, laterally to the vomer bone and ventrally to the cartilage of the nasal septum (Fig. 1B). The VNO is enveloped by a cartilaginous capsule (arrowheads), which, except in its dorsal part, completely surrounds the parenchyma of the organ. In its central part the vomeronasal ducts can be easily observed (Fig. 1D).

**Figure 1.**
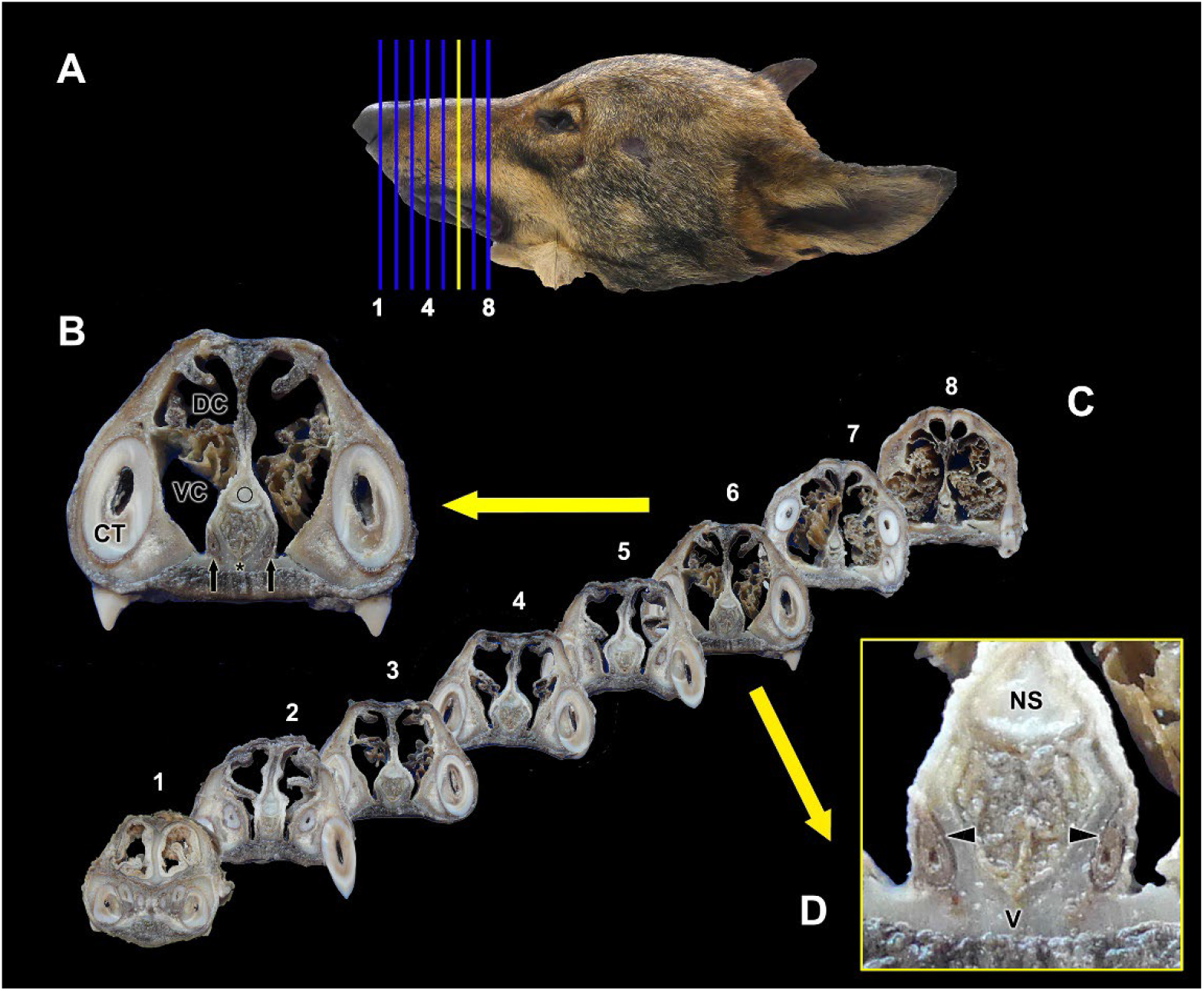
Macroscopic cross-sectional study of the nasal cavity of the wolf. **A.** Lateral view of the head of the wolf showing the eight consecutive levels chosen for the macroscopic sectional study. **B.** The central part of the vomeronasal organ (VNO) is located at level 6 (yellow line in A, enlarged section in B and D). The VNO corresponds to two tubular structures located in the ventral part of the nasal septum (black arrows), lateral to the vomer bone (asterisk), and ventral to the cartilage of the nasal septum (circle). **C.** Transverse sections of the nasal cavity ordered from rostral (1) to caudal (8), corresponding to the levels represented in (A). **D.** At higher magnification it can be seen how the VNO is enveloped by a cartilaginous capsule (arrowheads). In the central part of the VNO, the vomeronasal ducts can be observed. CT, canine tooth; DC, dorsal concha; NS, nasal septum; VC, ventral concha. Scale bars: B = 2 cm, D = 1 cm.

The study of the topographic relationships of the vomeronasal organ was extended to CT scans which allowed obtaining very clear images of the head cavities including the nasal concha, nasal meatuses, teeth, vomer bone and nasal septum. The bony configuration of the vomer bone and palatine fissure (Fig. 2A) deserved special attention, given the location of the VNO on the lateral surface of the vomer, as demonstrated by the previous sectional anatomical study of the nasal cavity. The obtained results are provided in three CT sections: horizontal (Fig. 2B), sagittal (Fig. 2C), and transverse (Fig. 2D). In all three planes, levels including the VNO have been chosen. The vomeronasal organ was placed bilaterally on each side of the vomer bone, at the most rostral and ventral region of the nasal cavity; a narrow compartment located dorsomedially to the palatine fissure and to the level of the root of the upper canine tooth.

**Figure 2.**
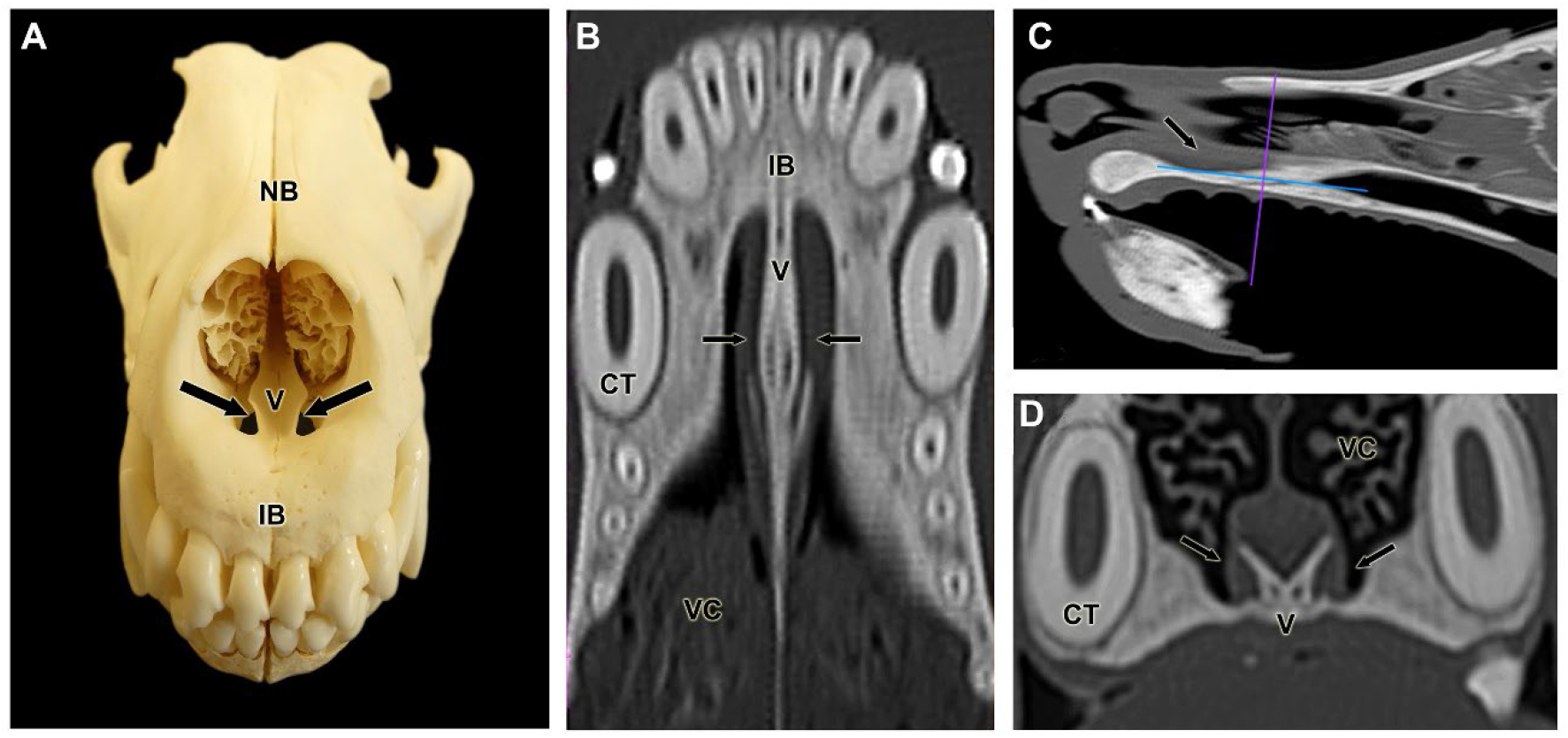
Topographical relationship of the wolf VNO. **A.** Fronto-dorsal view of the skull skeleton showing the relationship of the lateral part of the vomer bone (V) to the palatine fissure. **B-D.** Computerized tomography images of the head in the horizontal (B), parasagittal (C) and transverse (D) planes. Arrows indicate the location of the VNO. CT, canine tooth; IB: incisive bone; NB, nasal bone; VC, vomeronasal cartilage.

To access the VNO, it was necessary to expose the nasal septum. First, the lateral wall of the cavity formed by the maxillary bone was removed and then the large ventral nasal concha was extracted (Fig. 3). In the basal area of the septum, the VNO lies covered by respiratory mucosa layer. The presence of the caudal nasal myelinated nerve, which enters the VNO at its most caudal extremity (Fig. 3A), is a reliable indicator of the location of the VNO. On the mucosal surface of the nasal septum, amyelinic fibers of the vomeronasal nerve can be seen running in a caudo-dorsal direction toward the lamina cribrosa of the ethmoid. In their most rostral segment, the NVN are in direct relation to the respiratory mucosa and more caudally they are running into the olfactory mucosa, easily distinguished by their yellowish color (Fig. 3B). By removing the lateral wall of the cranial cavity, the size of the telencephalon and the location of the olfactory **z**bulb can be determined (Fig. 3C).

**Figure 3.**
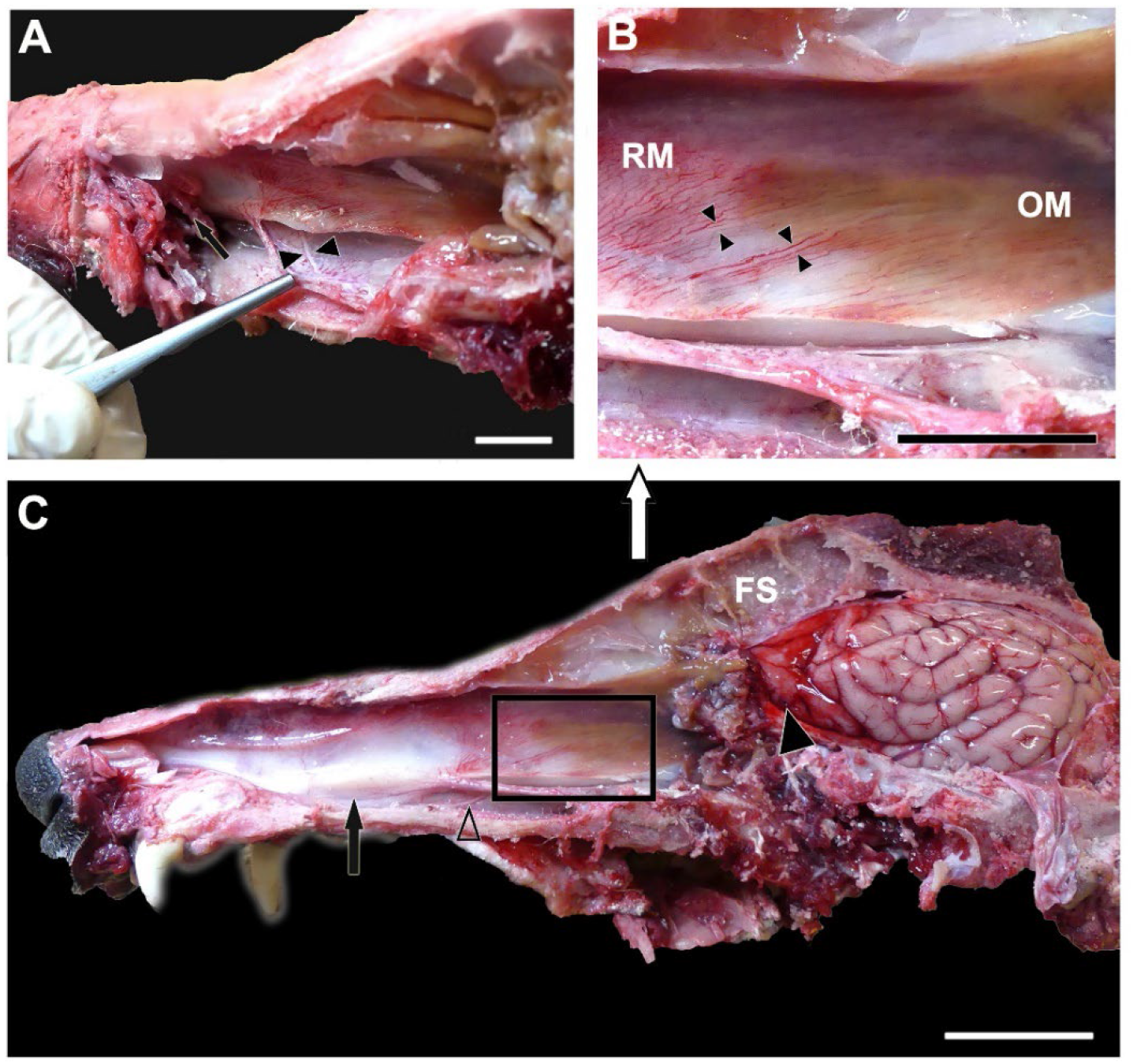
Dissection of the nasal and cranial cavities of the wolf. **A.** Dorsolateral view of the nasal cavity, showing the projection zone of the VNO (arrow) and the myelinated branches forming the caudal nasal nerve (arrowheads) entering the VNO. **B.** Enlargement of the inset shown in (C) displaying the vomeronasal nerves leaving the VNO (arrowheads) in a caudodorsal direction towards the medial portion of the cribriform plate of the ethmoid. RM: Respiratory mucosa. OM: Olfactory mucosa. **C.** Lateral view of the nasal and cranial cavities, showing in the latter the left-brain hemisphere. The projection area of the VNO (arrow), the vomeronasal nerves in a caudodorsal direction (inset), the caudal nasal nerve (open arrowhead) and the olfactory bulb (arrowhead) are also indicated. FS, Frontal sinus. Scale bar = 1.5 cm.

Prior to the removal of the VNO, the nasal cavity was reduced to small blocks containing the ventral part of the nasal septum and the bony floor of the palate. This made it possible to verify more clearly the relationship of the VNO with the nasal septum. At a caudal level the VNO is located on both sides of the base of the nasal septum, in a relatively elevated position due to the presence of a vertical projection of the vomer bone (Fig. 4A). At more rostral levels, the VNO is perfectly adapted to the lateral curvature of the vomer bone. In this case the VNO can be easily accessed from a ventral approach. Removing the respiratory mucosa from the nasal cavity that laterally overlies the organ allows the visualization of the shape and development of the organ (Fig. 4B). Finally, the VNO was dissected out from the bone tissue where it is held by dense connective tissue. To confirm the VNO identity, a transverse cut was made in the sample. The vomeronasal duct and the parenchyma of the organ are identifiable at first glance (Fig. 4C).

**Figure 4.**
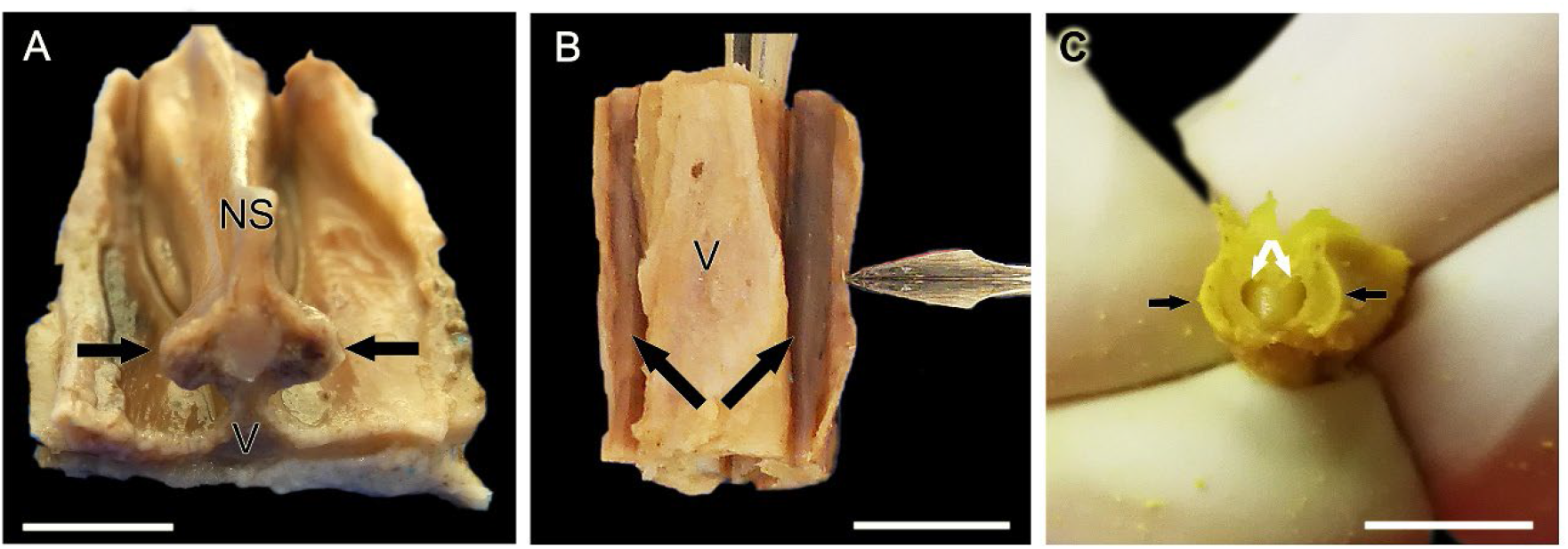
Wolf VNO after its full extraction in association to the vomer bone. **A.** Transverse cross section of the nasal septum corresponding to the level 9 shown in Figure 1. From a caudodorsal viewpoint, the VNOs (arrows) are visualized on both sides of the base of the nasal septum (NS) over the thin vertical projection of the vomer bone (V). **B.** On a more rostral level, corresponding to level 3 of Figure 1, a ventral view of the vomer bone show both VNOs with its distinctive rounded and elongated shape (arrows). The lancet points to the lifted respiratory mucosa of the nasal cavity that covers the VNO. **C.** Dissected out and partially transversely sectioned VNO (black arrows) showing the crescent shape vomeronasal duct (white arrows). A and B, fresh samples; C, Bouin’s fluid fixed sample. Scale bars: A-B = 0.5 cm, C = 0.2 cm.

To complete the anatomical study of the VNO, we examined how the VNO communicates with the external environment, which is necessary for the chemical messenger molecules to reach the neurosensory epithelium (Fig. 5). The wolf establishes this link indirectly, via the incisive duct (ID), which provides a passageway connecting the nasal and oral cavities (Fig. 5B), and the vomeronasal duct, which lays inside the middle of the two. The incisive papilla connects the incisive duct to the oral cavity (Fig. 5C). As it is not possible to establish macroscopically if the vomeronasal duct enters into the incisive duct, this will be evaluated microscopically.

**Figure 5.**
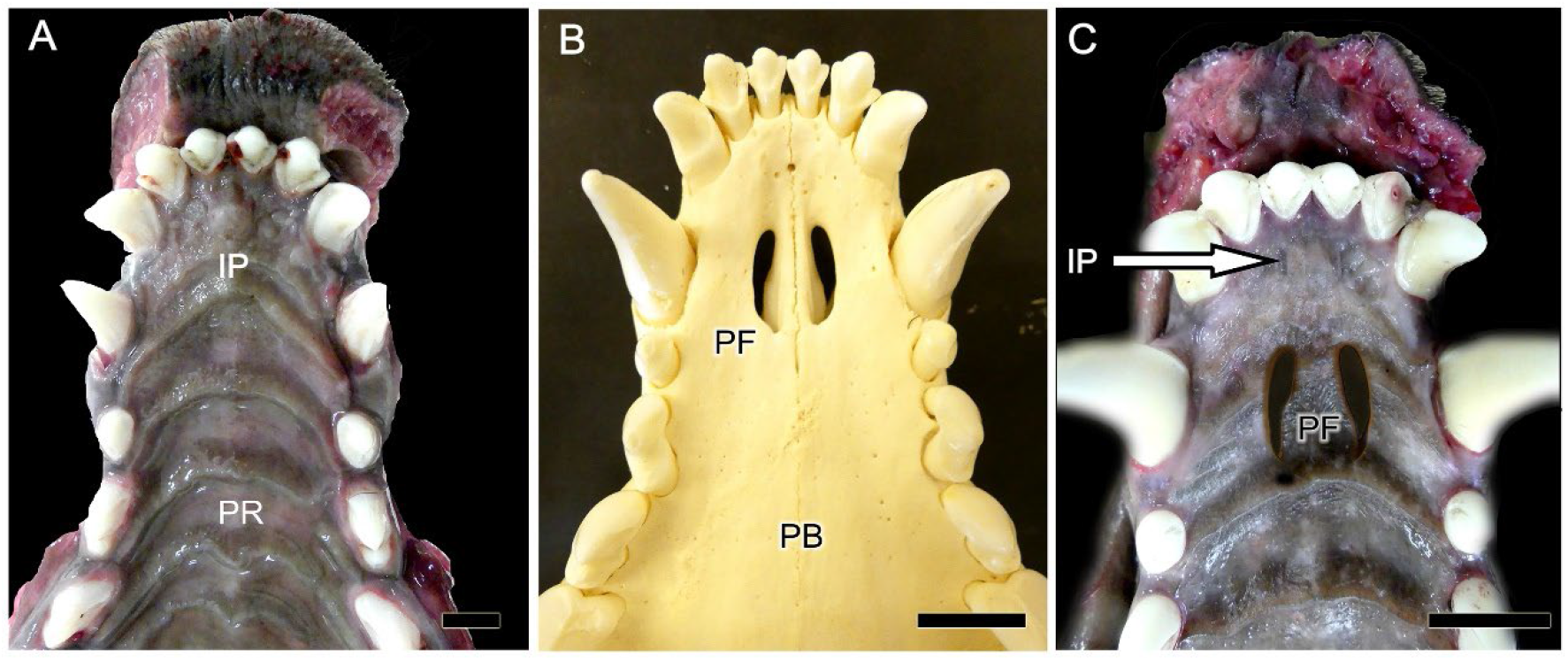
Ventral view of the palate of the wolf after removal of the mandible. **A.** The mucosa of the roof of the oral cavity is observed and in its most anterior part the location of the incisive papilla (IP) and palatine rugae (PR) are indicated. **B.** Skeleton of the corresponding area where the palatine fissures (PF) are observed on both sides of the midline. PB: Palatine bone. **C.** Superposition of images analogous to A and B to show the exact location of the PF on the mucosa of the palate. The IP is also shown (arrow). Scale bar: (A) 1 cm, (B-C) 1.5 cm.

#### 1.2 Accessory olfactory bulb (Fig. 6)

The macroscopic investigation of the brain of the wolf reveals that the olfactory bulbs are well developed and quite conspicuous in both the lateral and the medial view of the hemiencephalon (Fig. 6A-D). Particularly remarkable is the well-developed rhinencephalon, which possesses large olfactory pendunculi (Fig. 6E) and wide and convex piriform lobes (Fig. 6B). The main objective of the macroscopic study was to identify in situ the location of the accessory olfactory bulb. However, in none of the studied specimens we were able to carry out its identification due to its reduced dimerization and the usual presence of blood clots in the ethmoidal fossa. The anatomical tracing of the vomeronasal nerves in all cases pointed to an area located in the caudomedial part of the main olfactory bulb, which was the object of special attention in the sagittal histologic series of the bulb (Fig. 6G).

**Figure 6.**
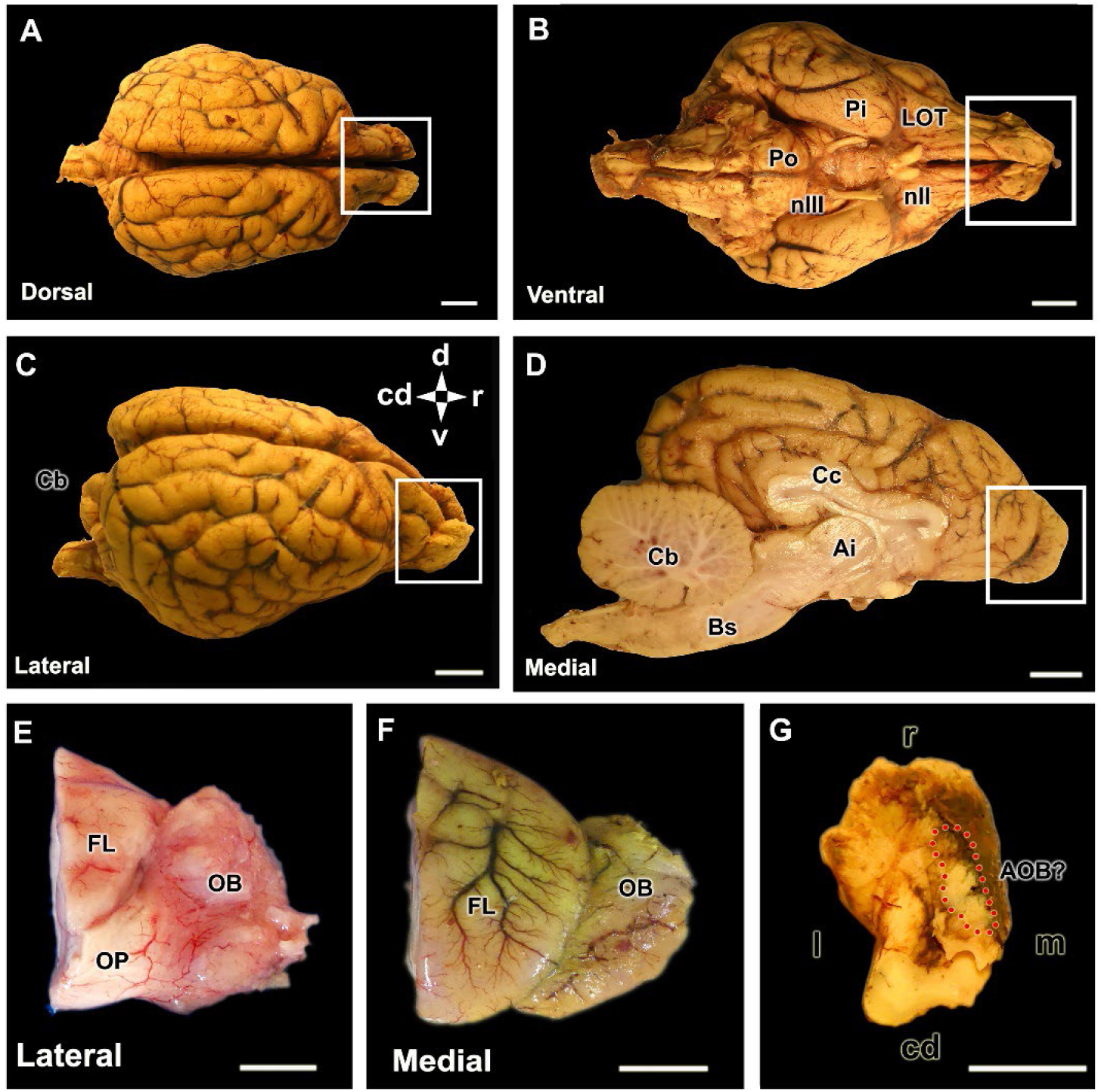
Encephalon and olfactory bulb of the wolf. **A-D.** Dorsal, ventral, lateral and medial views of a Bouińs fluid (BF) fixed encephalon. The white box highlights the dorsal, ventral, lateral and medial views of the olfactory bulb (OB), respectively. **E.** Lateral view of the right olfactory bulb, separated from a formalin-fixed brain. **F.** Medial view of the left MOB separated from the BF-fixed brain. **G.** Dorsocaudal view of the left OB, fixed in BF. The area where the accessory olfactory bulb is presumptively located is indicated by the broken red circle. Bs, Brain stem; Cb, Cerebellum; Cc, Corpus Callosum; FL, Frontal lobe; Ia, Interthalamic adhesion; LOT, lateral olfactory tract; nII, Optical nerve; nIII, Oculomotor nerve; OP, olfactory peduncle; Pi, piriform lobe; Po, Pons; R, rostral. Cd, caudal; d, dorsal; l, lateral; m, medial. Scale bar = 1 cm.

### 2. Microscopic study of the VNO (Fig. 7-14)

As a first microscopic approximation, the VNO consists of a vomeronasal capsule and a vomeronasal duct, which is surrounded by parenchyma, which contains blood vessels, nerves, and vomeronasal glands (Fig. 7). The capsule encloses the parenchyma and prevents the lumen from collapsing. It has “U” shape and is comprised of hyaline cartilage. The lumen is typically elliptical and lined by pseudostratified epithelium. The lateral portion of this epithelium is of a respiratory nature, while the medial portion is of a neurosensory type, and both portions appear to be of comparable size and development.

**Figure 7.**
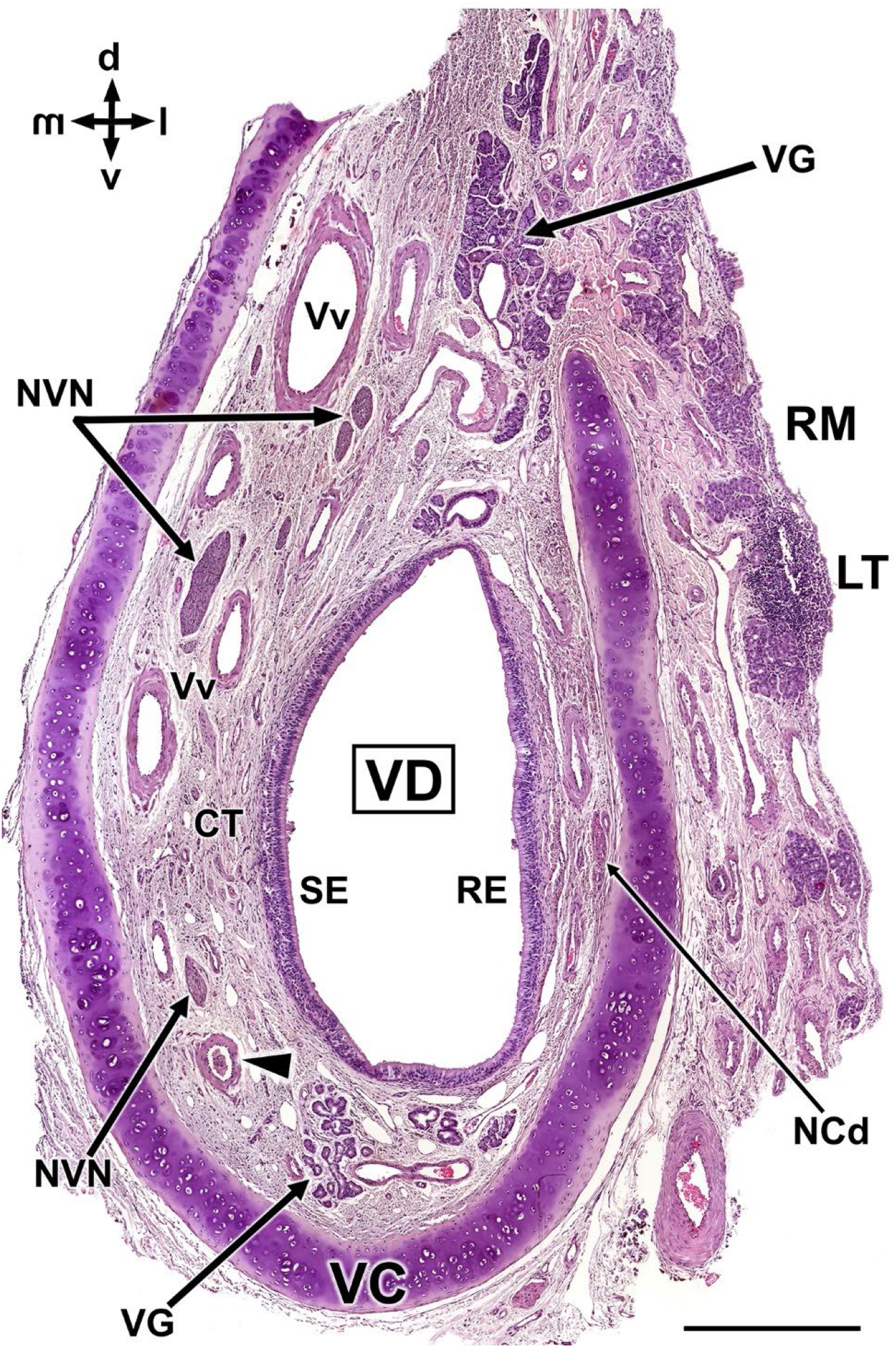
Histological transverse section of the wolf vomeronasal organ in its central portion, stained with haematoxylin-eosin. This central level exemplifies the major histological features of the VNO. VC: vomeronasal cartilage. In its central portion, it forms an incomplete ring which opens dorsally. VD: vomeronasal duct. NCd: caudal nasal nerve. SE: neurosensory epithelium. RE: respiratory epithelium. NVN: vomeronasal nerves. RM: respiratory mucosa. VG: vomeronasal glands. LT: Lymphoid tissue. CT: connective tissue. Vv: vomeronasal veins. Arrowhead: artery. D, dorsal; m, medial; l, lateral; v, ventral. Scale bar = 500 μm.

The blood vessels are the major component of the parenchyma of the organ, distributed throughout its circumference, although they are predominant in the dorsal and lateral part where numerous large and muscular veins can be seen. There are hardly any arteries and these have a reduced size. Numerous branches of the vomeronasal nerves are located between the medial veins. In the lateral parenchyma the nerves are of smaller caliber and correspond to branches of the caudal nasal nerve. The glandular component is not very abundant, concentrating the glands near the ventral and especially dorsal commissures. The dorsal glands invade the parenchyma of the respiratory mucosa, which also has a profuse irrigation, abundant glandular tissue and diffuse lymphoid tissue. The parenchyma of the VNO is completed with connective tissue, very abundant throughout the whole length of the organ.

The histological study was extended to sections located in the rostral and caudal third of the VNO (Fig. 8). In both cases the general organization of the organ is similar. In the anterior third (Fig. 8A) the cartilage is more open dorsally, the venous vessels have a larger caliber but thinner wall and the nerve branches are less abundant. The differentiation between both epithelia is visible. Gallego’s trichrome staining shows the remarkable specific development of the connective tissue. In a most caudal section (Fig. 8B), the medial branch of the cartilage increases its development, producing a “J” shape. At this level, the vomeronasal nerves leave the parenchyma by their dorsal extremity forming large nerve trunks.

**Figure 8.**
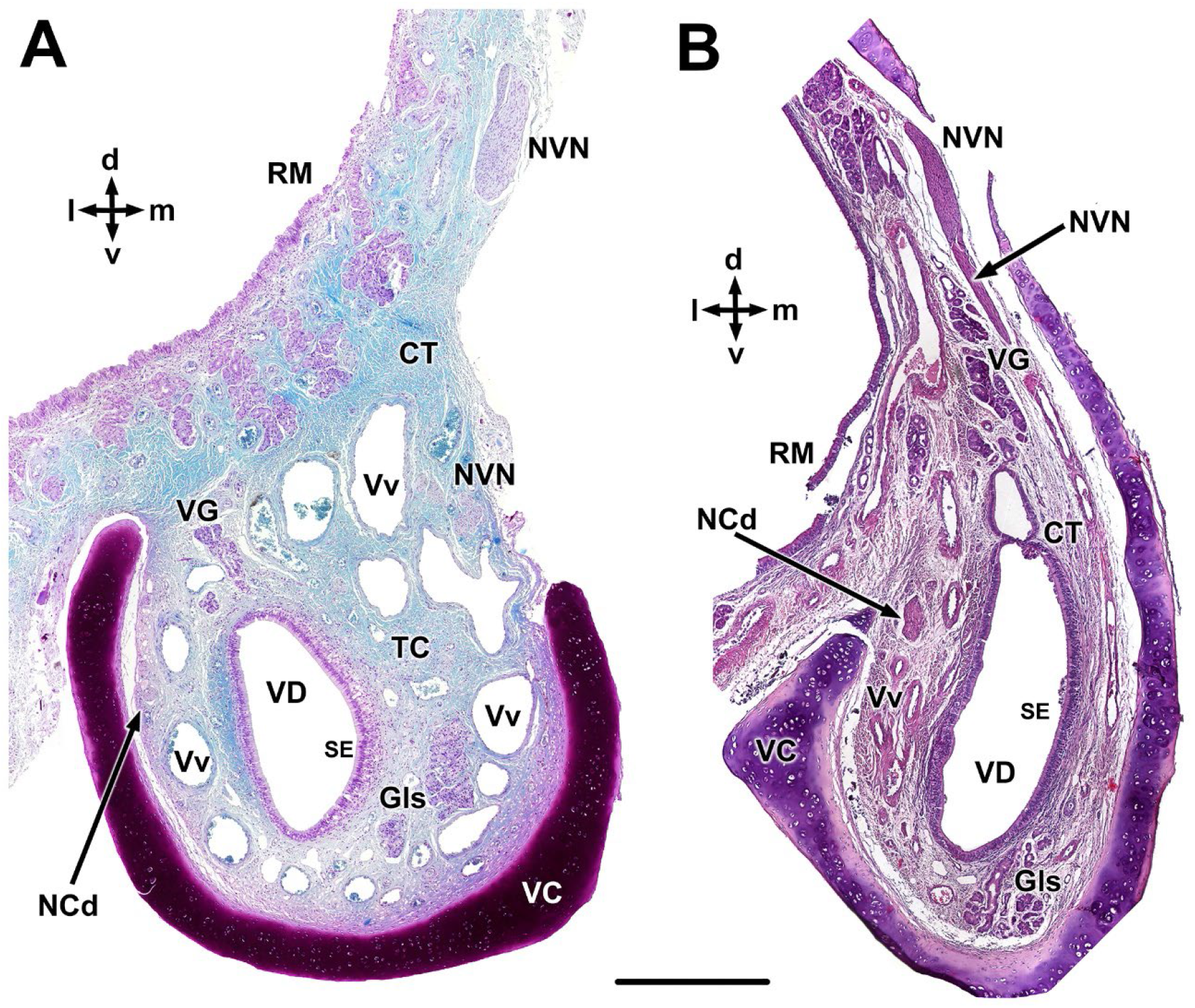
Transverse sections of the wolf VNO at two selected levels. A. Rostral level. **B.** Caudal level. VC, Vomeronasal cartilage; VD, vomeronasal duct; NCd, caudal nasal nerve; SE, Sensory epithelium; NVN, vomeronasal nerve; RM, respiratory mucosa; VG, vomeronasal glands; CT, connective tissue; Vv, vomeronasal veins. Staining: Gallego’s trichrome (A) and hematoxylin-eosin (B). d, dorsal; l, lateral; m, medial; v, ventral. Scale bar = 500 μm.

The histological study of the two epithelia lining the vomeronasal duct (Fig. 9) was performed with Gallego’s trichrome (Fig. 9A-C, F-G) and hematoxylin-eosin stains (Fig. 9D,E). The broad lamina propria associated with the neuroreceptor epithelium contains abundant connective tissue, amidst its fibers are situated vessels and branches of the vomeronasal nerves (Fig. 9A). The most striking fact is the presence in a subepithelial position in the lamina propria of conspicuous clusters of tightly packed cells that traverse the basal cell layer, maintaining a direct relationship with the neuoreceptor cells (Fig. 9B). Although the cells forming the clusters lack visible processes their nuclei are similar in shape, content, and staining to those of the neuroreceptor cells. To our knowledge this is the only example of a species presenting such cellular organization in the vomeronasal epithelium. More superficial to them are the cellular components of the vomeronasal neuroepithelium (Fig. 9C).

**Figure 9.**
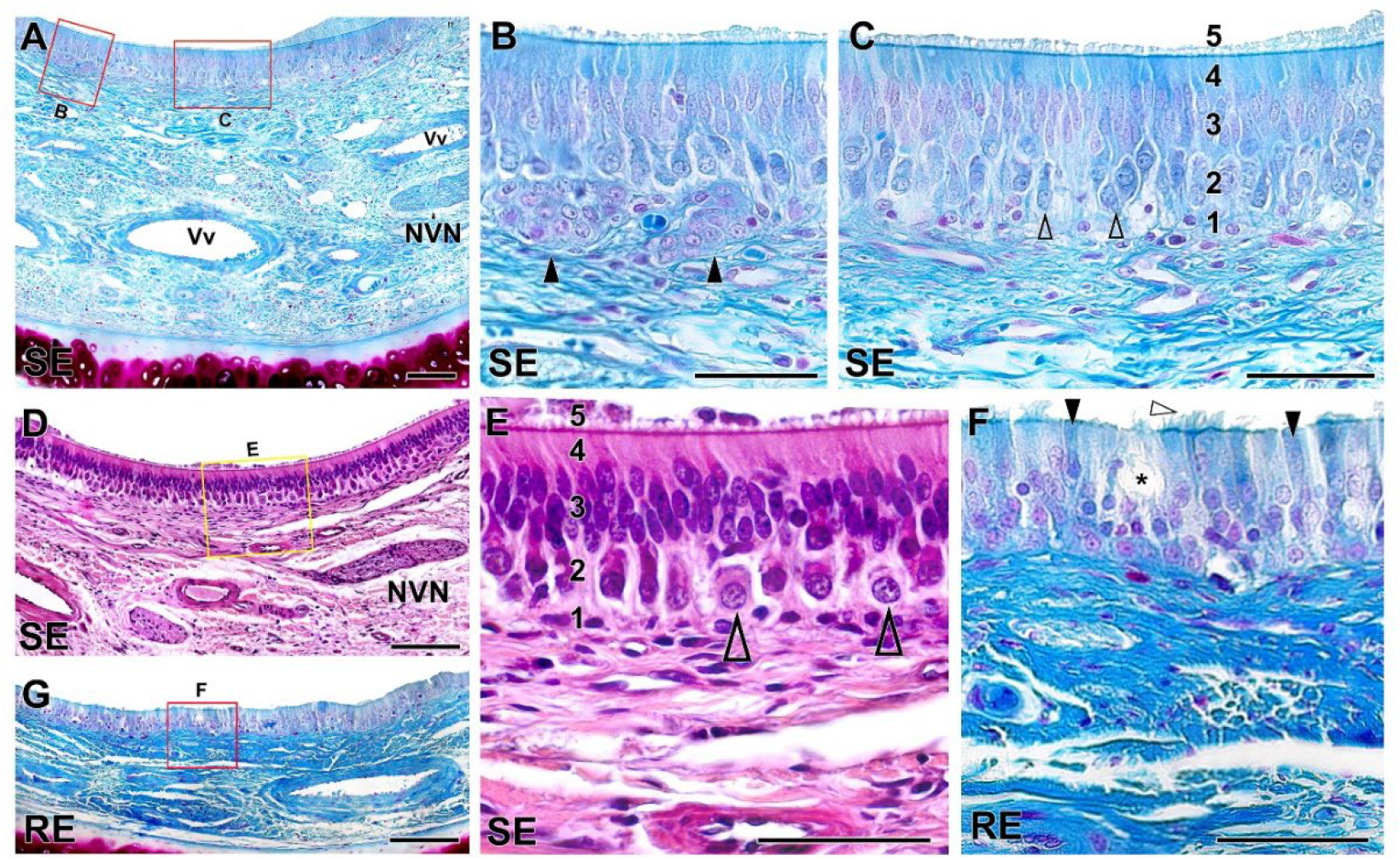
Histological study of the epithelia of the wolf vomeronasal duct. **A.** Medial parenchyma of the organ stained with Gallego’s trichrome. The overlying sensory neuroepithelium (SE), enlarged in (B,C), shows a large lamina propria with veins (Vv), vomeronasal nerves (NVN) and connective tissue. **B.** Two large clusters of cells similar in appearance to neuroreceptor cells (arrowheads) are observed in the lamina propria. **C.** Cellular components of the neuroepithelium: 1, basal cells; 2, neuroreceptor cells (open arrowheads); 3, sustentacular cells; 4, cell processes layer; 5, mucomicrovilliar complex. **D.** Neurosensory epithelium and lamina propria, stained with haematoxylin-eosin. **E.** Enlargement of the area shown in D showing the five components of the sensory neuroepithelium. Open arrowheads: neurosensory cells. **F.** Enlargement of the area of respiratory epithelium (RE) shown in (G), stained with Gallego’s trichrome. Note the presence of cilia (white arrowhead), chemosensory cells (black arrowheads) and goblet cells (*). Scale bars: A, D, G = 100 μm; B, C, E, F = 50 μm.

They are organized in a columnar, pseudostratified, and non-ciliated epithelium. From basal to luminal its components are basal cells, neuroreceptor cells, sustentacular cells, the cell processes layer, and the superficial mucomicrovilliar complex. The neuroreceptor cells are ellipsoidal in shape and are not densely packed, which allows visualization of their fine axonal and dendritic processes. Their nucleus is rounded and has a visible nucleolus. The sustentacular cells have densely packed nuclei in the apical position and are elongated in shape. Hematoxylin-eosin staining confirmed these observations (Fig. 9D,E). The respiratory epithelium (Fig. 9F,G) is a pseudostratified columnar epithelium that presents cilia on its surface. It is composed by sustentacular, chemosensory and goblet cells (Fig. 9F).

The VNO parenchyma of the wolf presents numerous glands, which are more developed in its dorsal area (Fig. 10). Both Gallego’s trichrome (Fig. 10A) and hematoxylin-eosin (Fig. 10B) stainings show the serous tubuloalveolar nature of the vomeronasal glands. Alcian blue staining shows that the acini secrete an Alcian Blue-positive secretion (Fig. 10C).

**Figure 10.**
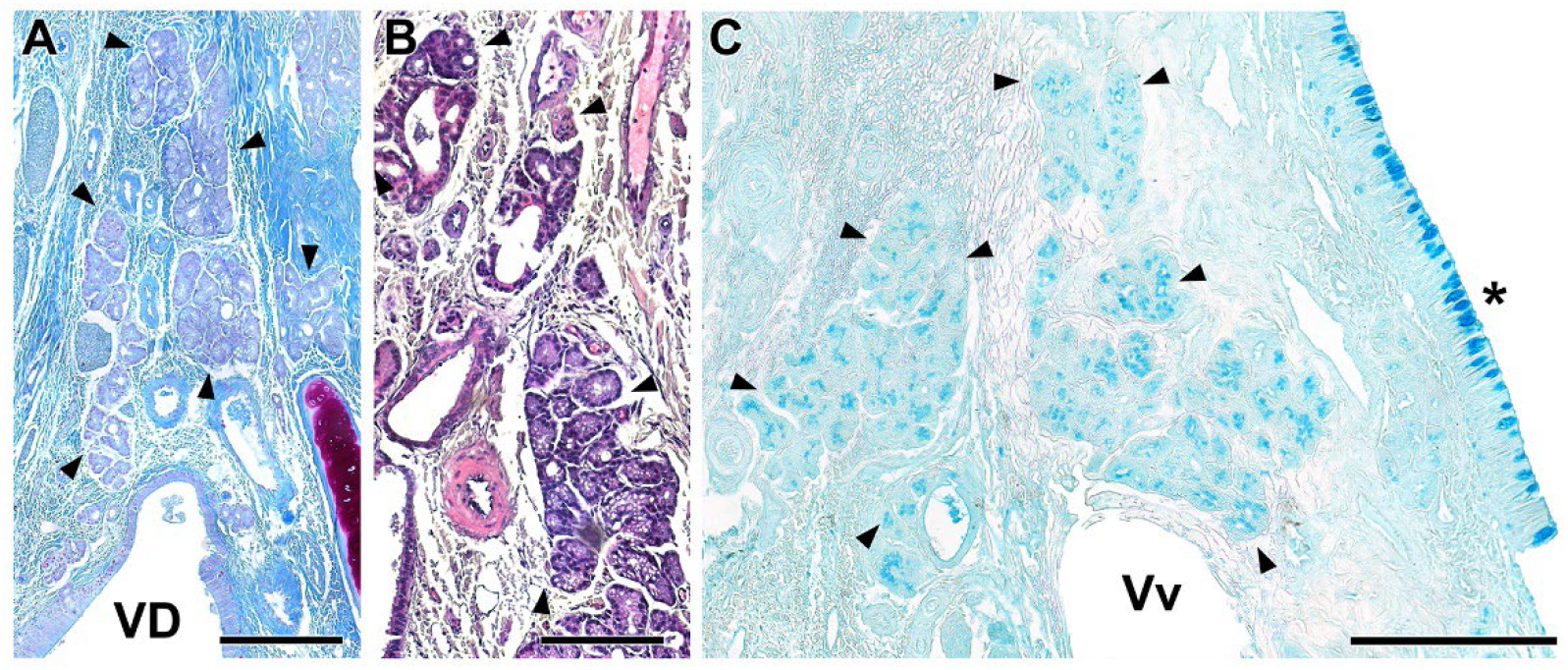
Vomeronasal glands of the wolf. The VNO parenchyma presents an abundant number of glands, which are more developed in its dorsal area. **A-B.** The arrowheads delimit the glandular areas of interest. Both Gallego’s trichrome (A) and hematoxylin-eosin (B) stainings show the serous tubuloalveolar nature of the wolf vomeronasal glands. **C.** Alcian blue staining shows the positivity of these acini to this stain. VD, vomeronasal duct; Vv, vomeronasal veins; (*) Respiratory mucosa. Scale bar = 250 μm.

The information regarding the blood vessels is summarised in Fig. 11. These vessels form a large vascular network, providing the organ with erectile tissue functionality. In the anterior portion of the VNO, veins form a complete vascular ring surrounding the vomeronasal duct (Fig. 11A). In the central area of the VNO, large venous sinuses run along the lateral portion of the parenchyma (Fig. 11D). Conversely, in the caudal area of the VNO where the glove-fingered termination of the vomeronasal duct appears, numerous medium sized veins predominate (Fig. 11C). In contrast, arteries throughout the VNO are small and sparse (Fig. 11B,C).

**Figure 11.**
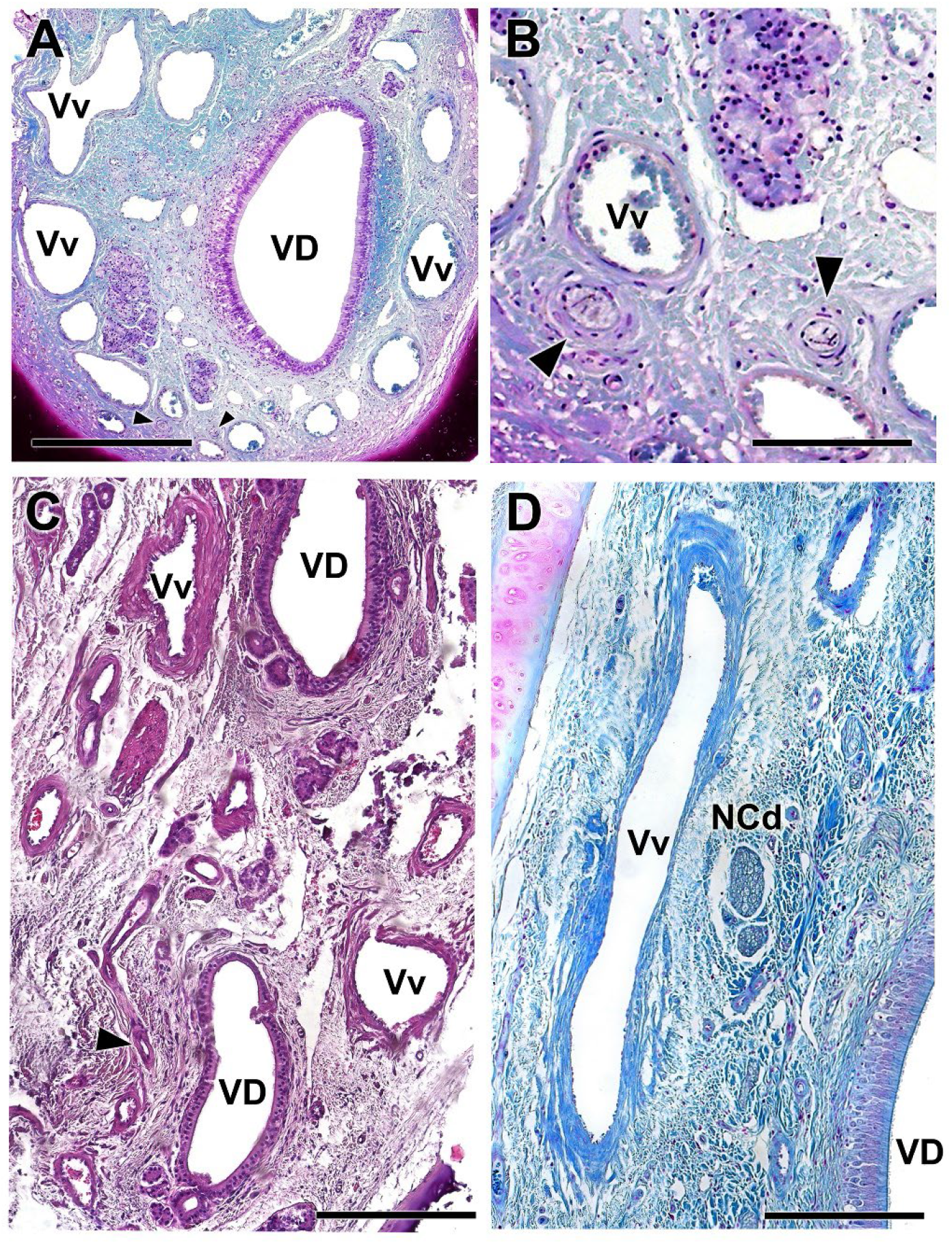
Vasculature of the wolf VNO. **A.** Image of the anterior portion of the VNO shown in Fig. 8A exemplifies the presence of a profuse venous ring (Vv) surrounding the vomeronasal duct (VD). The arteries (arrowheads) are small and sparse. **B.** Arteries indicated in A showed at higher magnification. **C.** Caudal section of the VNO showing the glove-fingered termination of the vomeronasal duct (VD). Numerous veins (Vv) and small arterial trunks (arrowhead) predominate at this level. **D.** In a central section of the VNO, large venous sinuses (Vv) run along the lateral portion of the parenchyma. Staining: (A,B,D) Gallego’s trichrome; (C) hematoxylin-eosin. Scale bars: A, D = 250 μm; C = 100 μm; B = 50 μm.

The innervation of the VNO consists of two types of nerves classified according to their myelination (Fig. 12). The sensory component is comprised of unmyelinated branches of the vomeronasal nerves (Fig. 7, 8). These nerves occupy the medial parenchyma of the VNO, characterised by their homogeneous appearance, and densely packed nerve bundles (Fig. 12B). The lateral part of the parenchyma contains myelinated nerve branches, originating from the caudal nasal nerve (Fig. 7, 8B, 11D). These branches have a looser structure, and the empty spaces corresponding to the myelin sheaths are visible (Fig. 12A).

**Figure 12.**
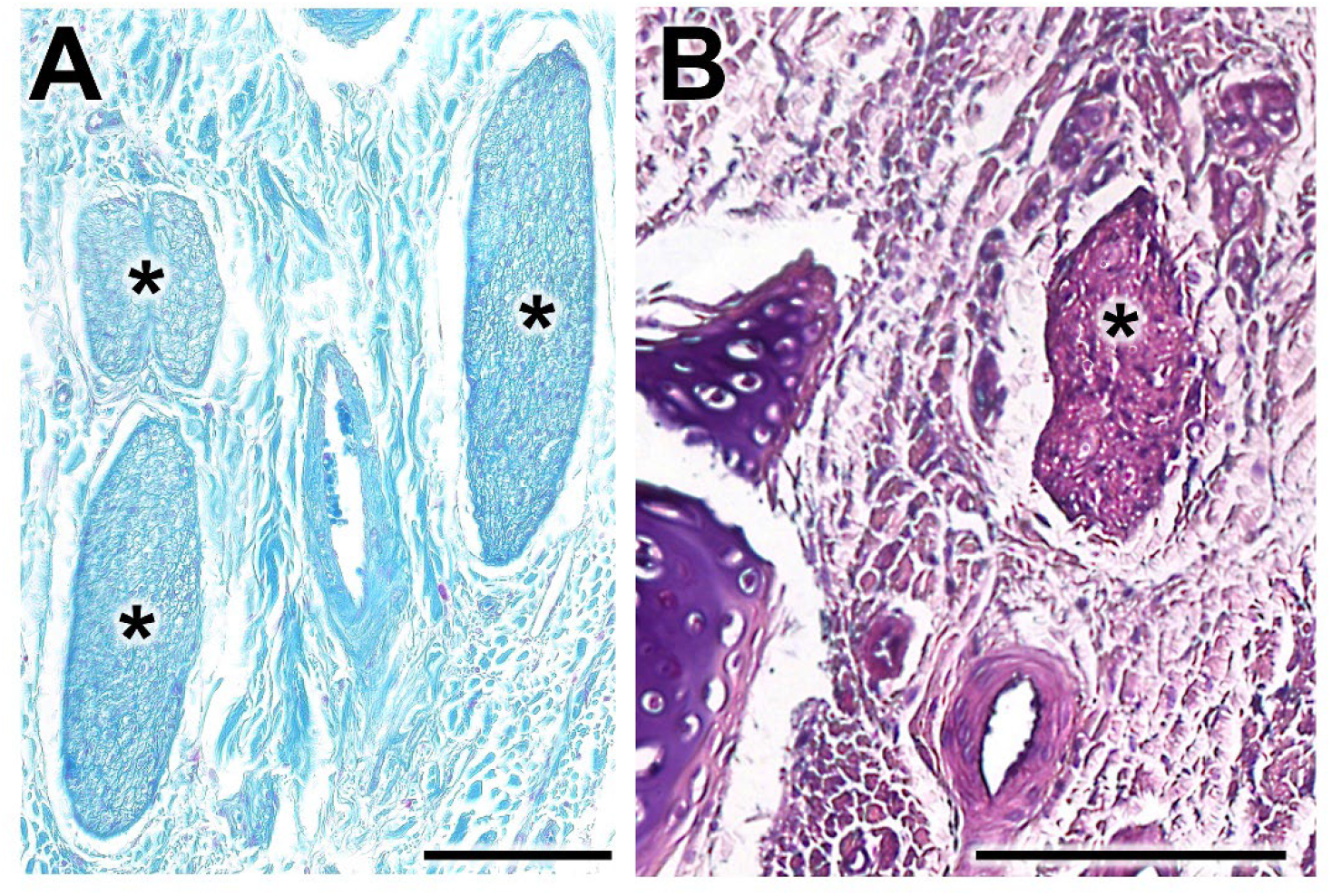
Histological study of the wolf vomeronasal organ innervation. **A.** Unmyelinated branches of the vomeronasal nerves (*). They are characterised by their homogeneous appearance, with densely packed nerve bundles. **B.** The lateral part of the parenchyma contains myelinated nerve branches originating from the caudal nasal nerve (*). Their structure is looser and the void spaces corresponding to the myelin sheaths are visible. Staining: A, Gallego’s trichrome; B, hematoxylin-eosin. Scale bar = 100 μm.

Our microscopic study of the VNO was supplemented with an examination of decalcified histological sections from the rostral part of the nasal septum. These sections illustrate the topographical relationship of the rostral VNO with the vomer bone (Fig. 13). At this level, the vomeronasal cartilage presents an elongated morphology, accompanying the shape of the vomer bone, and laterally it exhibits a wide gap. Examination of more rostral decalcified samples allowed confirmed microscopically the functional communication of the vomeronasal duct with the external environment via the incisive duct. The latter runs laterally to the VNO, facilitating direct communication of the vomeronasal duct with both the oral and nasal cavities, as the incisive duct opens ventrally into the aperture present in the elevated mucosa, forming the incisive papilla, and dorsally into the meatus present in the floor of the nasal cavity (Fig. 14).

**Figure 13.**
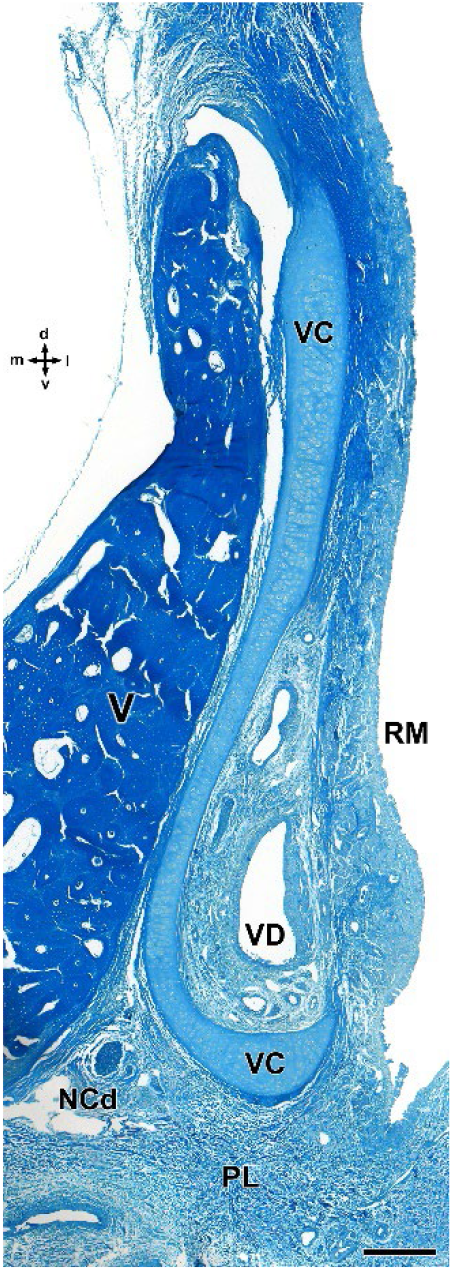
Decalcified cross-section of the anterior portion of the wolf nasal septum, stained with Gallego’s trichrome, showing the topographical relationship of the VNO with the vomer bone. The vomeronasal cartilage (VC) presents an elongated morphology, accompanying the shape of the vomer bone (V), and opens laterally. LP, lamina propria; NCd, caudal nasal nerve; PL, palate; RM, respiratory mucosa; VD: vomeronasal duct. d, dorsal; l, lateral; m, medial; v, ventral. Scale bar = 250 μm.

**Figure 14.**
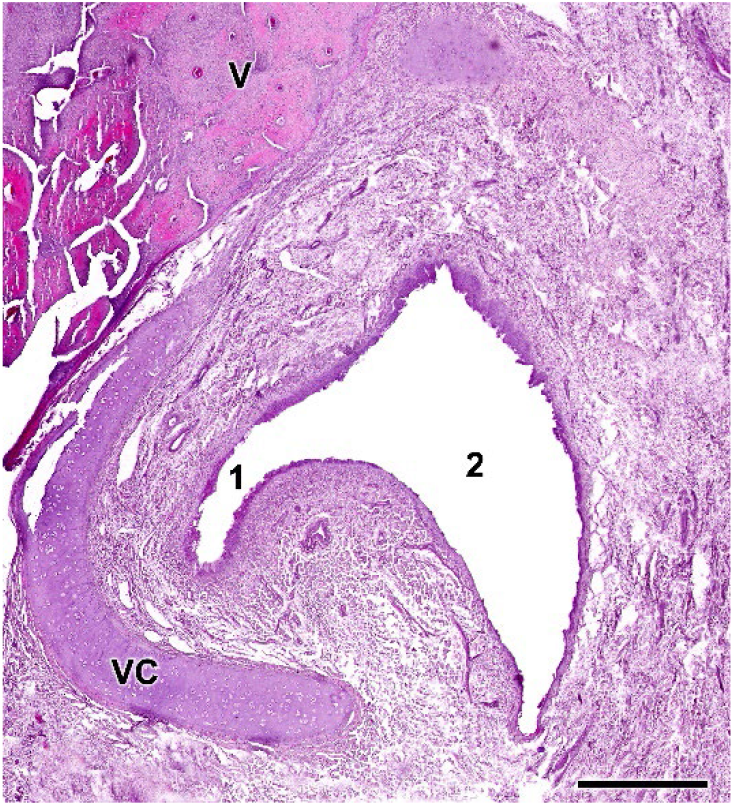
Opening of the vomeronasal duct into the incisive duct of the wolf. The vomeronasal duct (1) establishes a direct communication with the incisive duct (2). Anatomically, the latter runs laterally to the organ, allowing a double direct communication between the oral and nasal cavities. Staining: hematoxylin-eosin. VC, vomeronasal cartilage; V, vomer bone. Scale bar = 500 μm.

### 3. Lectin histochemical study of the VNO (Fig. 15)

Both the neuroreceptor cells of the sensory epithelium and the vomeronasal nerves of the VNO show histochemically positive labelling for both UEA and LEA lectins (Fig. 15). However, when considering the neuroepithelial cell clusters located in a basal position in the neuroepithelium, both lectins exhibit contrasting labelling patterns.

**Figure 15.**
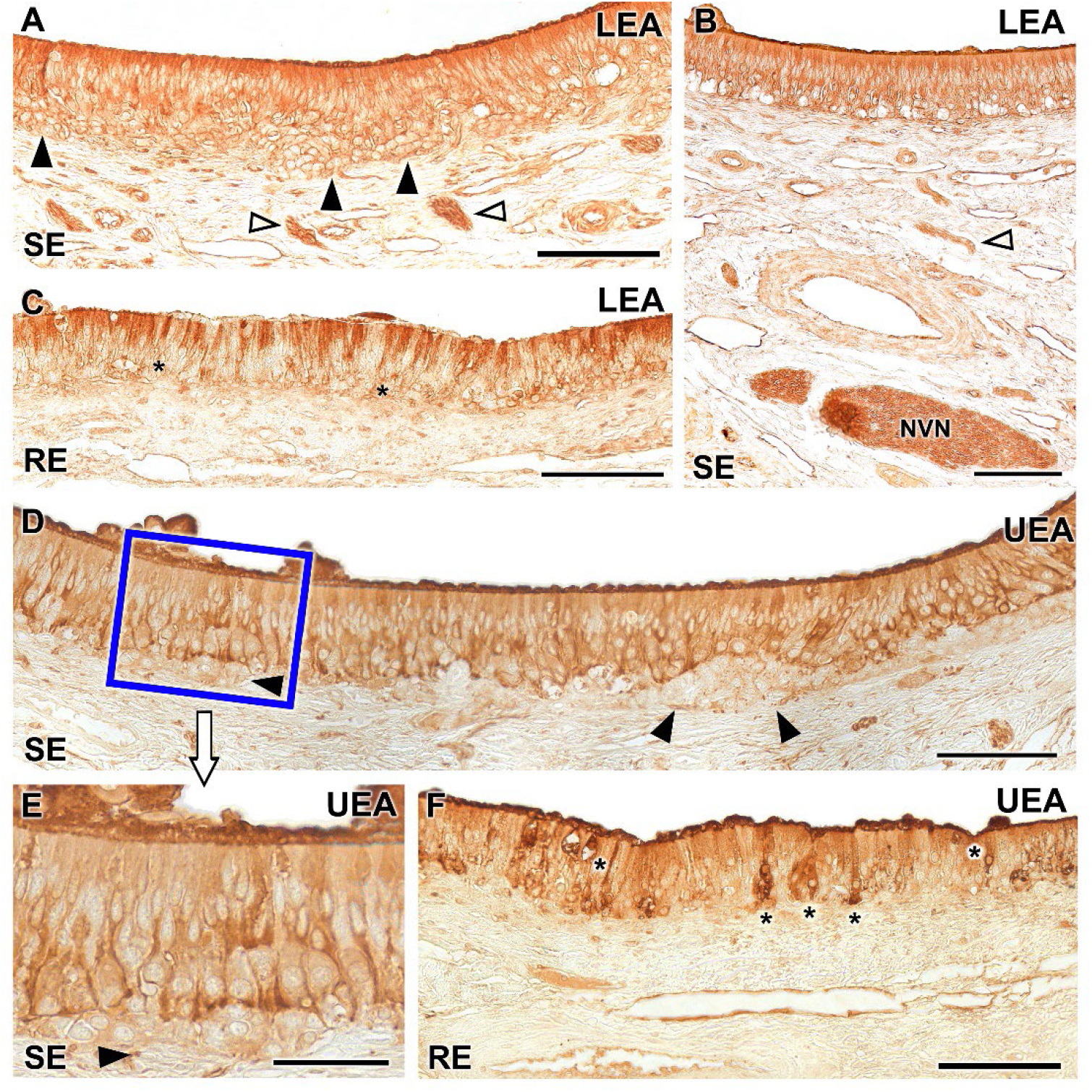
Lectin histochemical labelling of the vomeronasal epithelium. **A-C.** LEA lectin labelling: The neuroreceptor epithelium (SE) shows intense labelling throughout all its components including neuroepithelial clusters (black arrowheads) and nerve bundles in the lamina propria (white arrowheads) (A). This labelling extends to the vomeronasal nerves (NVN) (B). **C.** The respiratory epithelium shows a diffuse LEA positivity, concentrated mainly in the most apical part of the epithelium, while the basal cells remain unlabelled (asterisk). **D-F.** UEA lectin: Positivity is demonstrated across the sensory epithelium of the VNO excluding the neuronal clusters (arrowheads) which are not labelled (D). At higher magnification, an intensity gradient increasing with depth can be discerned (E). The unlabelled neuronal clusters are shown (arrowhead). **F.** The respiratory epithelium exhibits fainter labelling, with few positive cells scattered along the epithelium (*). UEA labelling is also markedly concentrated in the mucociliary layer. Scale bar: A-D, F = 100 μm; E = 50 μm.

While LEA produces positive labelling of neuronal clusters (Fig. 15A), UEA lectin does not label these cells (Fig. 15. D, E). LEA labelling is slightly more intense in the apical processes of the epithelium (Fig. 15A, B), while UEA produces more intense labelling in the basal areas of the epithelium (Fig. 15D, E), excluding the unlabelled clusters. Regarding the respiratory epithelium, it exhibits diffuse labelling from both lectins, albeit with distinct patterns. LEA positivity is primarily concentrated in most of the apical processes of the epithelium (Fig. 15C). UEA presents a few strongly labelled cells scattered along the epithelium. In the mucociliary complex, the labelling is stronger with UEA than with LEA (Fig. 15F).

### 4. Immunohistochemical study of the VNO (Fig. 16)

The anti-Gαo antibody, which specifically labels the αo subunit of the G-protein transduction cascade, associated to the V2R receptor, labelled a subpopulation of neurons mostly present in the basal layers of the neuroepithelium which extend their axons to the adjacent vomeronasal nerves (Fig. 16A, B). The labelling comprises the entire length of the immunopositive neuroreceptor cells, from the apical dendrite to the soma. Immunopositive neuroreceptor cells embrace the intraepithelial capillaries of the VNO (Fig. 16C). The anti-Gαi2 antibody, which labels the αi2 subunit of the G-protein transduction cascade associated to the V1R receptors, labels neuroreceptor cells present in the central zone of the epithelium and the vomeronasal nerves (Fig. 16D, E). Unlike Gαo, no immunopositive neurons are identified around the intraepithelial capillaries. The dendritic knobs are less numerous than those labelled by anti-Gαo but more dilated (Fig. 16E). The deep neuronal clusters are immunonegative.

**Figure 16.**
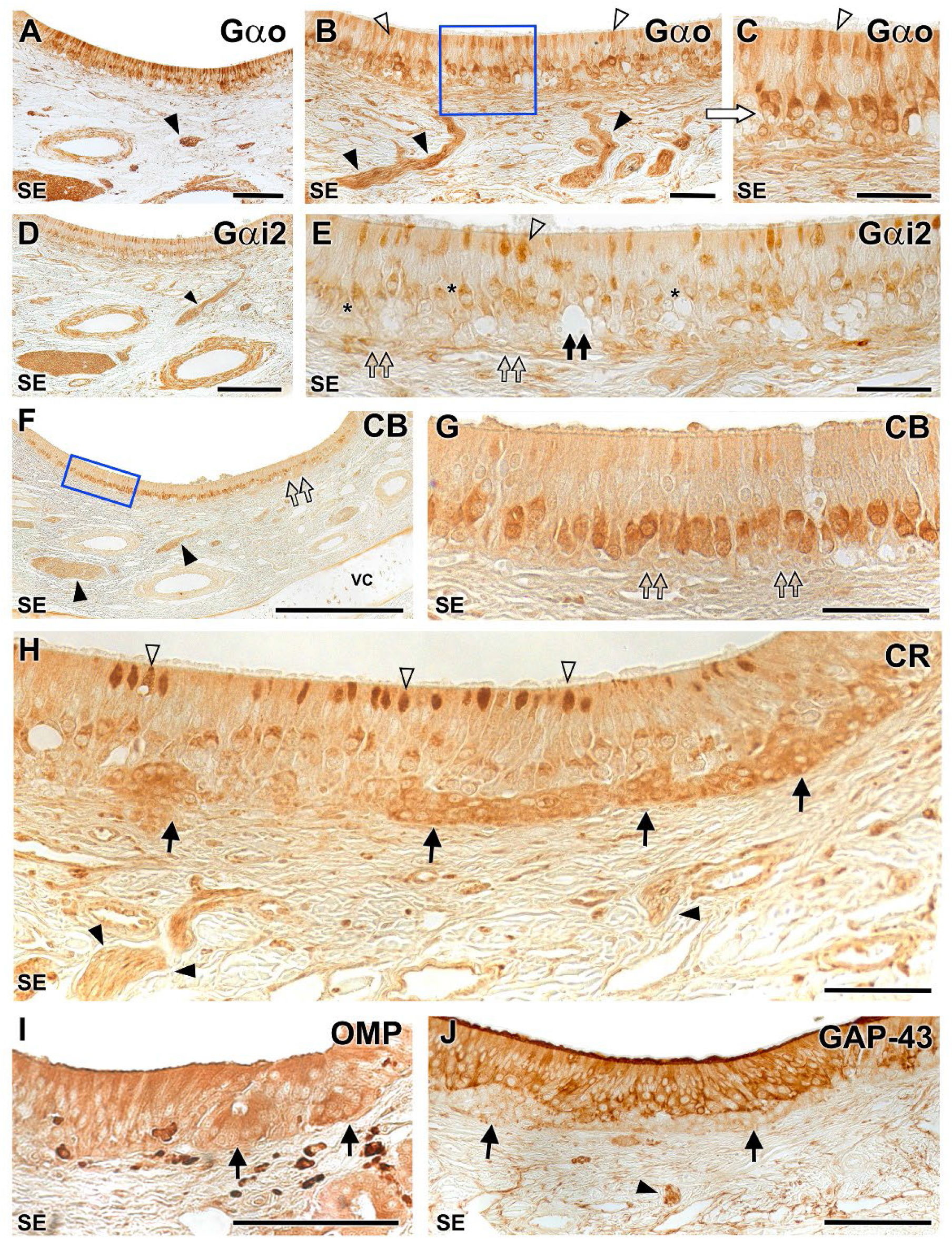
Immunohistochemical study of the wolf VNO. **Gαo (A-C)**: Immunolabelling with anti-Gαo shows a pattern of neuronal labelling concentrated in the neuroreceptor cells present in the basal layers of the neuroepithelium (A, B) and extending to the adjacent vomeronasal nerves (arrowheads). At higher magnification (C: enlarged area of the blue box in image B), it is appreciated how the labelling extends along the entire length of the immunopositive neuroreceptor cells, from the apical dendrite (open arrowhead) to the soma. Immunopositive neuroreceptor cells embrace the intraepithelial capillaries of the VNO. **Gαi2 (D-E)**: The labelling is concentrated on the neuroreceptor cells present in the central zone of the epithelium (asterisk) and the vomeronasal nerves (arrowhead). Unlike Gαo, no immunopositive neurons are identified around the intraepithelial capillaries (black double arrow). The dendritic knobs are less numerous than in Gαo but more dilated (open arrowhead). The deep neuronal clusters are immunonegative (open double arrows). **Calbindin (CB) (F-G)**: At low magnification (F) uniform labelling is observed throughout the neuroreceptor cell layer, extending into the vomeronasal nerves (black arrowheads). No immunopositivity is observed in the clusters (double arrows). At higher magnification (G: magnification of the blue box in F) is appreciated how the immunopositive cells correspond to a regularly aligned subpopulation in an intermediate zone between the basal and apical layers. The terminal knobs are poorly labelled. **Calretinin (CR) (H)**: A strong immunopositivity to the neuronal clusters is observed (arrows). In addition, a subpopulation of neuroreceptor cells whose dendrites show dilated terminal knobs can be identified (white arrowheads). Vomeronasal nerves (black arrowhead). **OMP (I)**: Anti-OMP produces a diffuse labelling throughout the epithelium, including the basal clusters (arrows). **GAP-43 (J)**: Pattern similar to (I) but without labelling the neuronal clusters (arrows). Arrowhead: vomeronasal nerves. SE: Neurosensory epithelium. VC: Vomeronasal cartilage. Scale bars: A, D, I, J = 100 μm; B, C, E, G = 250 μm; H = 50 μm.

Anti-calbindin produces a uniform immunolabelling throughout the neuroreceptor cell layer, extending which also comprises the vomeronasal nerves (Fig. 16F). The immunopositive cells correspond to a regularly aligned subpopulation in an intermediate zone between the basal and apical layers. The terminal knobs are poorly labelled and no immunopositivity is observed in the basal clusters (Fig. 16G). Anti-calretinin shows a strong immunopositivity in the basal neuronal clusters (Fig. 16H)). In addition, a subpopulation of neuroreceptor cells whose dendrites show dilated terminal knobs can be identified. Vomeronasal nerves are also stained.

Anti-OMP, which binds to OMP, a protein that acts as a marker of neuronal maturation, produces a diffuse labelling throughout the epithelium, including the basal clusters (Fig. 16I) whereas anti-GAP-43, which binds to GAP43, a protein associated with neuronal axonal growth, produces a similar pattern to but without labelling the neuronal clusters (Fig. 16J).

### 5. Histological study of the nasal septum mucosa (Fig. 17)

To follow the pathway of the vomeronasal nerves through the nasal septum, after dissecting out nasal mucosa an immunohistochemical labelling was carried out using antibodies against the G alpha subunits proteins (Fig. 17).

**Figure 17.**
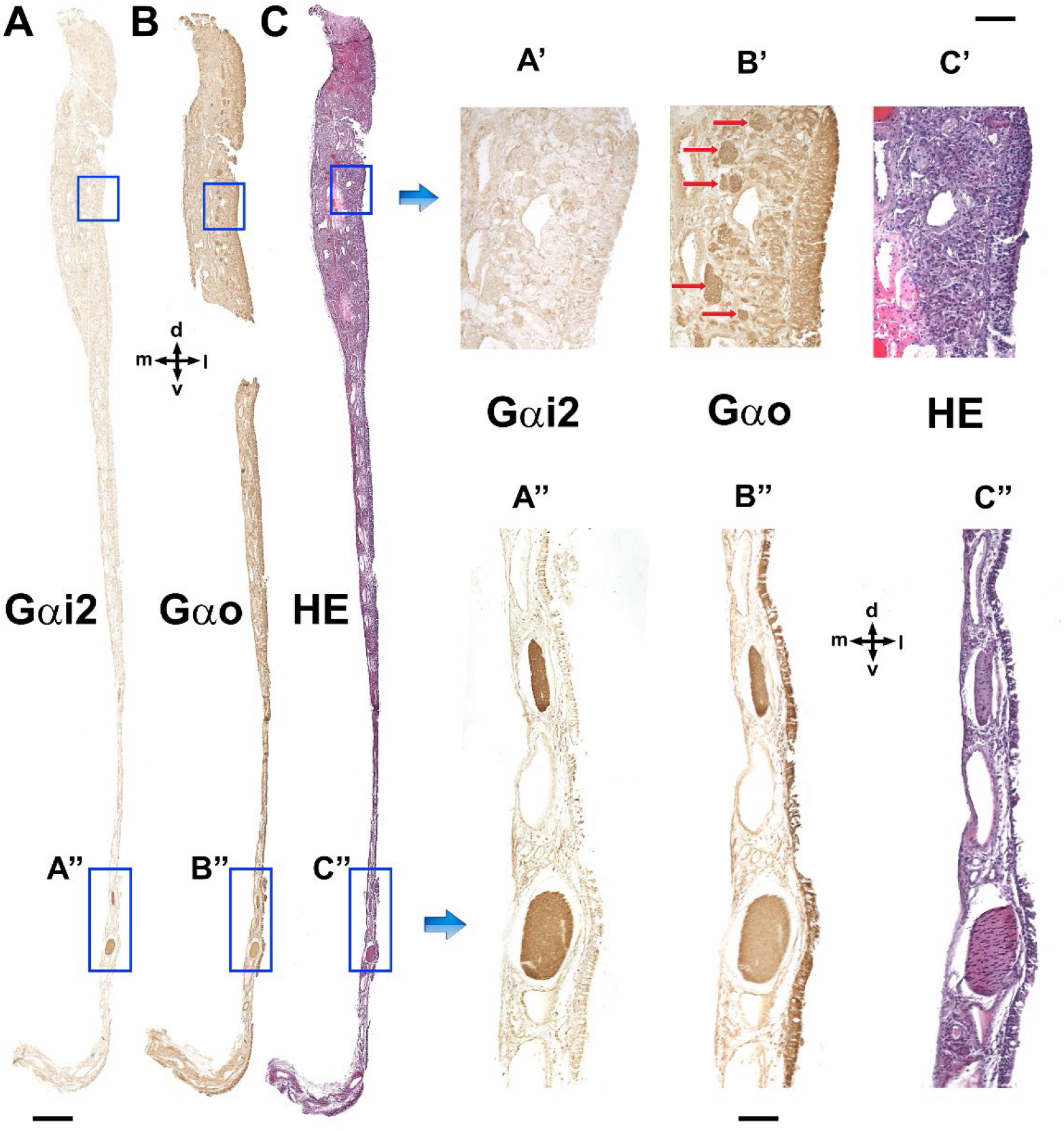
Immunolabelling of the VNO nasal septum mucosa with anti-G proteins subunits. **A.** Anti-Gαi2 exclusively stains the vomeronasal nerves as they run through the mucosa of the nasal septum (A’, A‘’). **B.** Anti-Gαo (B) stains both the vomeronasal nerves (B’’) and the olfactory nerves in the mucosa (B’, red arrows). **C.** Hematoxylin-eosin adjacent section. d, Dorsal; l, lateral; m, medial; v, ventral. Scale bars: A-C = 500 μm; A’-C’ and A’’-C’’ = 100 μm.

Whereas the Gαi2 subunit produced a clear and specific labelling of the vomeronasal nerves (Fig. 17A, A’, A”), the antibody against Gαo not only stained the vomeronasal nerves but also the olfactory nerves that course through the mucosa of the nasal cavity and the olfactory neuroepithelium that gives rise to these axons (Fig. 17B, B’, B”).

### 6. Histological study of the AOB (Fig. 18)

The histological structure of the accessory olfactory bulb was examined using hematoxylin and eosin (Fig. 18A,B, D, E) staining and Nissl staining (Fig. 18C,F-H), both performed on serial sagittal sections. Both stains revealed a significant development of this structure, particularly in relation to its two superficial layers: the nerve layer and the glomerular layer (Fig. 18A, D-F, H). The nerve layer represents the arrival point of the vomeronasal nerves (Fig. 18B, C), while the glomerular layer consists of well-defined, broad, and rounded glomeruli that are clearly visible with both stains (Fig. 18B, C). The mitral cells are diffusely distributed in a wide zone located between the glomerular layer and the granular layer, thus precluding the distinction between true plexiform and mitral layers. Therefore, the term ’mitral plexiform layer’ MPL) is used to describe this region (Fig. 18G, H). The granular layer displays clusters of small, rounded cells interspersed within the white matter (Fig. 18G).

**Figure 18.**
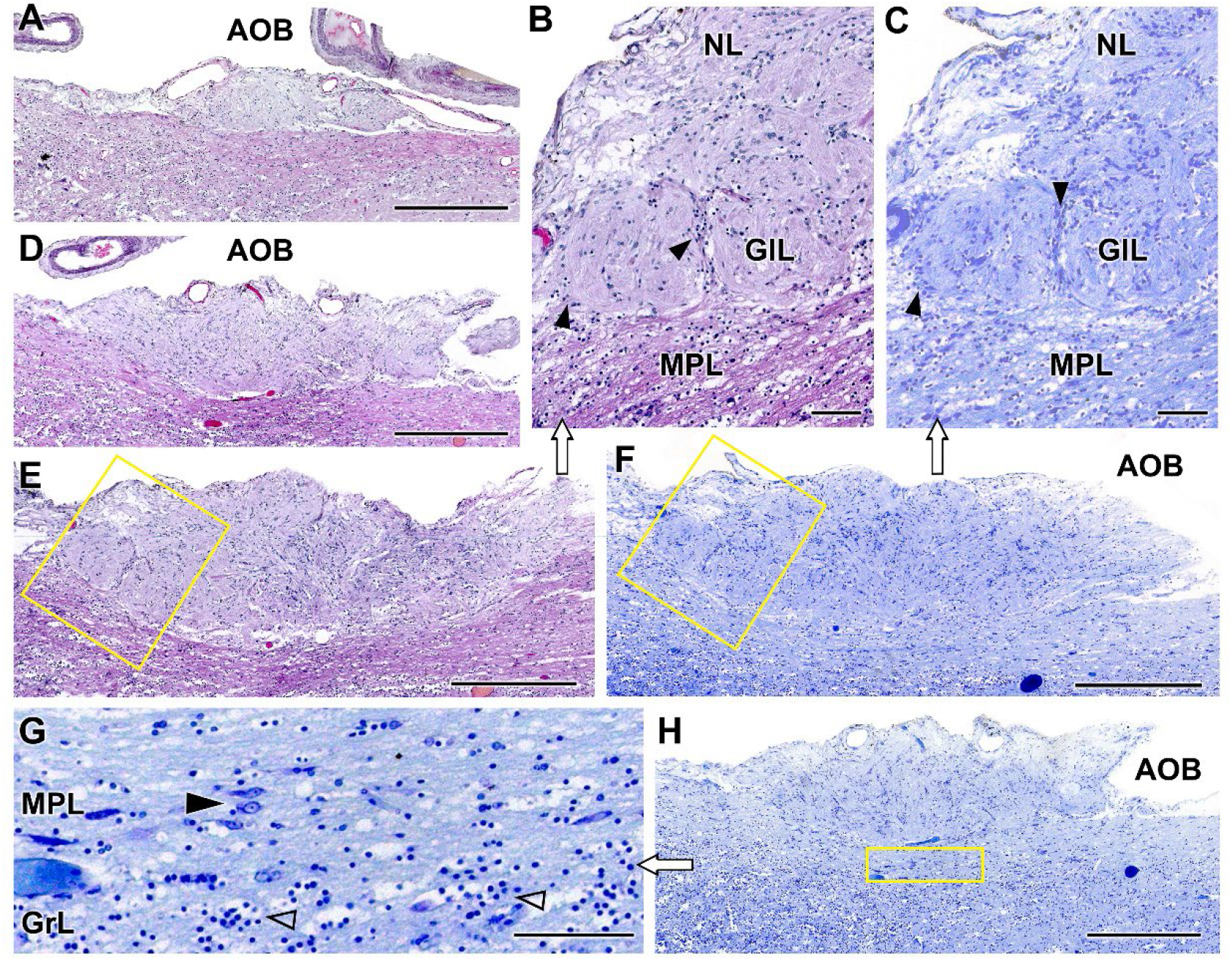
Histological study of the wolf accessory olfactory bulb stained with haematoxylin-eosin and Nissl staining. **A, D, E.** A general view of the AOB can be seen at three selected sagittal levels stained with HE. They show the elongated shape of this structure and the predominance of the nervous (NL) and glomerular layers (GlL). **B.** At higher magnification (corresponding to box in E) two glomerular formations clearly defined by periglomerular cells (arrowheads) are appreciated. **F, H**. Sagittal Nissl-stained sections at low magnification allows to appreciate the development of the AOB. **C.** The magnification of the superficial area of the AOB (box in F) allows to discriminate the presence of a mitral-plexiform layer (MPL). G. Enlargement of the deep area of the AOB (corresponding to box in H) shows mitral cells (black arrowhead) in the MPL as well as granular cells (open arrowhead) in the deeper granular layer (GrL). Scale bars: A, D, E, F, H = 500 μm; B, C, G = 100 μm.

### 7. Immunohistochemical study of the AOB (Fig. 19)

The immunohistochemical study of wolf AOB with G-protein subunits antibodies produced a complementary labelling for both Gαi2 and Gαo subunits. Whereas anti-Gαi2 immunostained uniformly and intensely the superficial nervous and glomerular layers of the AOB layers (Fig. 19A) anti-Gαo produced a reverse pattern, with the neuropil surrounding the AOB superficial layers being strongly immunopositive. Both the mitral-plexiform and granular layers of the AOB were inmunostained with anti-Gαo. However, the superficial layers were immunonegative although some immunopositive punctae areas were observed (Fig. 19B). This immunolabelling corresponded to the dendritic projections of mitral cells within the glomerular layer. The calcium binding proteins, calbindin (Fig. 19C) and calretinin (Fig. 19D), as well as OMP (Fig. 19E) showed an identical labelling pattern to that obtained with anti-Gαi2, with the immunolabelling concentrated in both the superficial layers, contrasting with an immunonegative neuropil.

**Figure 19.**
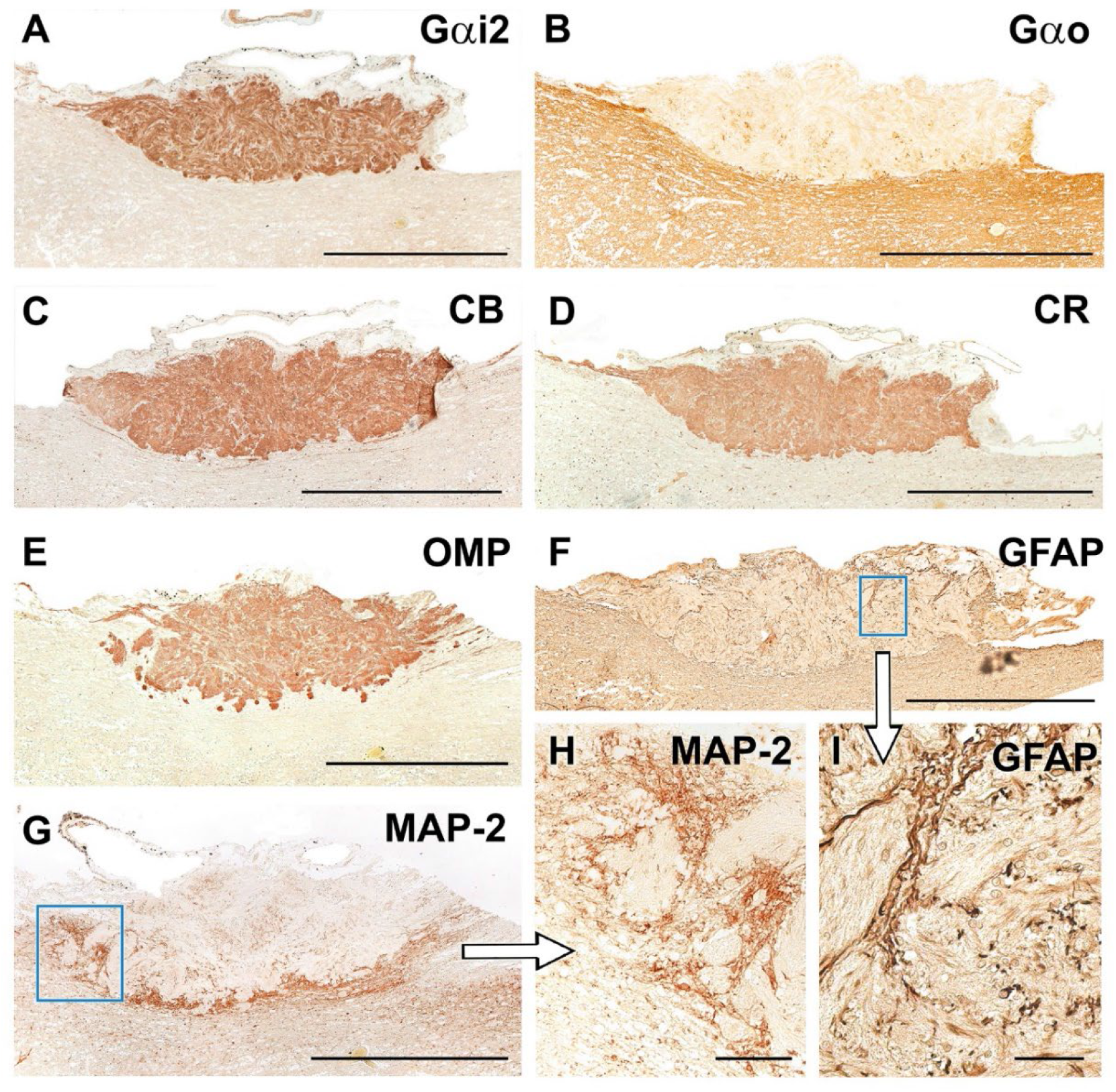
Immunohistochemical study of wolf AOB. **A.** Anti-Gαi2 uniformly and intensely label the superficial layers of the AOB (nervous and glomerular layers). The entire surrounding neuropil is negative. **B.** Anti-Gαo produces a reverse pattern to the one shown in A, where the neuropil surrounding the superficial layers is strongly immunopositive, including the mitral-plexiform and granular layers of the AOB. However, the superficial layer is clearly negative although immunopositive punctae areas are observed. **C-E.** The calcium binding proteins, calbindin (C) and calretinin (D), as well as OMP (E) show an identical labelling pattern to that obtained with anti-Gαi2, concentrated in both the nerve and glomerular layers and immunonegative for the neuropil. **F.** Anti-GFAP (F and enlarged area in I) produces a trabecular labelling pattern in the nerve and glomerular layers, which corresponds to the ensheathing glia accompanying the vomeronasal nerve endings. Occasionally, cell bodies belonging to these glial cells are visible. **G.** Anti-MAP-2 immunolabelling does not produce immunopositive labelling in the superficial layers (nervous and glomerular), but it strongly labels an irregular band corresponding to the MPL layer. **H.** MAP-2 immunopositive prolongations originating from the MPL can be observed running between the glomeruli of the AOB. (H: enlargement of the box in G). Scale bars: A-G = 500 μm; H = 50 μm.

Anti-GFAP, specific marker of glial cells, produced a trabecular labelling pattern in the nerve and glomerular layers, which corresponded to the ensheathing glia accompanying the vomeronasal nerve endings (Fig. 19F). Occasionally, cell bodies belonging to these glial cells were visible (Fig. 19I). Anti-MAP2 is a good marker for the somata and dendritic projections of the principal cells of the olfactory bulb. It strongly labelled an irregular band corresponding to the MPL layer (Fig. 19G). MAP-2 immunopositive prolongations originating from the MPL could be observed running between the glomeruli of the AOB (Fig. 19H).

### 8. Lectin histochemical study of the AOB (Fig. 20)

Both lectins employed labelled the AOB. UEA lectin selectively labelled the superficial layers of the AOB (Fig. 20A). The entire surrounding neuropil is negative. LEA lectin produced a similar labelling to UEA with a strong labelling in the AOB superficial layers, which prevented to differentiate both the nervous and glomerular layers (Fig. 20B). The surrounding neuropil showed a diffuse labelling pattern.

**Figure 20.**
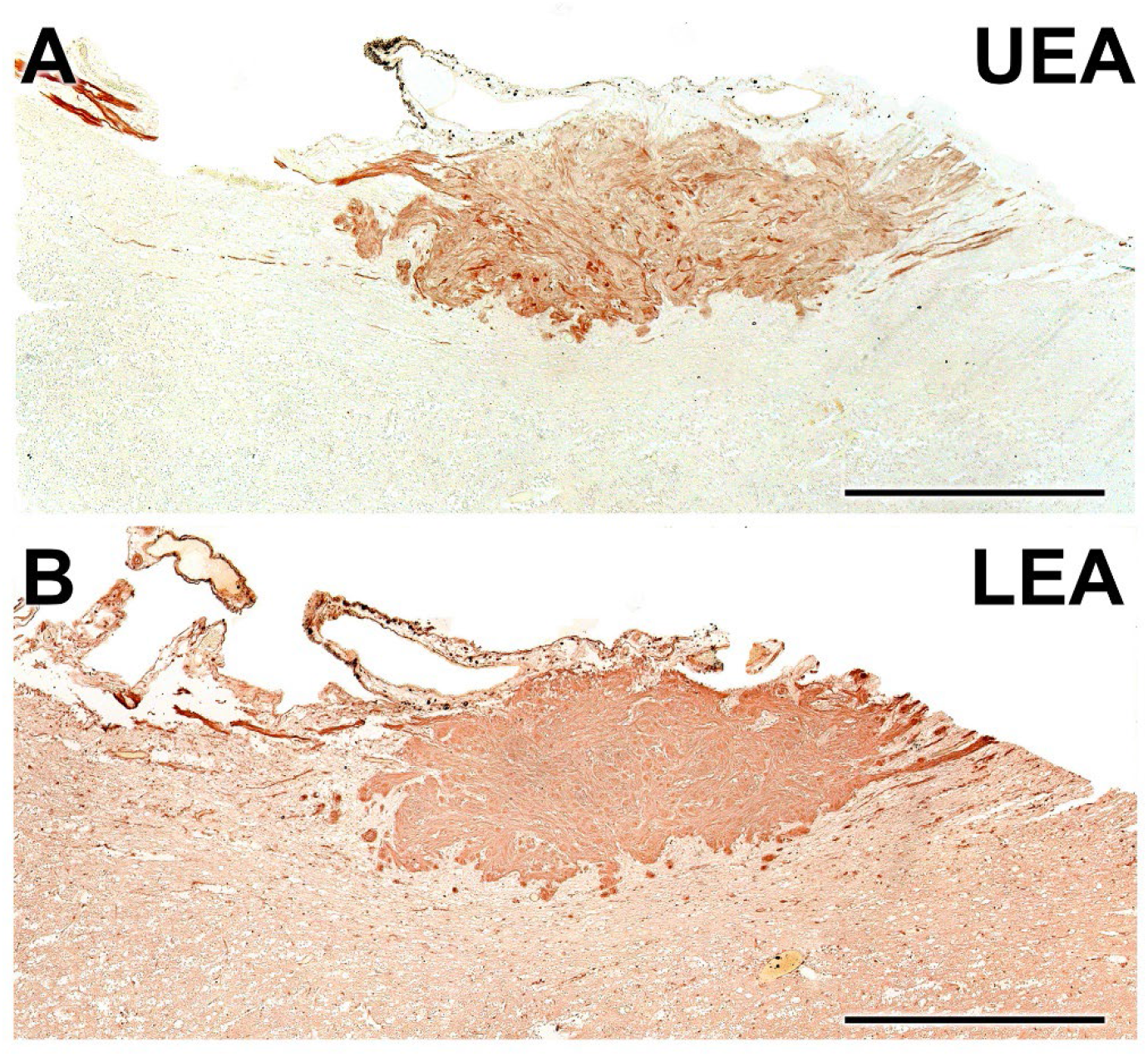
Lectin-histochemical study of the wolf AOB. **A.** UEA lectin labels uniformly and strongly the superficial layers of the AOB (nervous and glomerular layers). The entire surrounding neuropil is negative. **B.** LEA-lectin produces a similar labelling to UEA but with higher intensity in the AOB superficial layers and a diffuse pattern in the surrounding neuropil. Scale bar = 500 μm.

## DISCUSSION

Chemical communication, facilitated by pheromones, has long been recognized as an essential component of social and sexual interactions among canids. These complex chemical signals, detected and processed by the vomeronasal system, serve an array of functions ranging from mate selection and social hierarchy establishment to territory marking and are integral to the reproductive physiology of these animals (Gorman and Trowbridge 1989).

In this study, we have aimed to contribute to the ongoing study of the neuroanatomical and neurochemical aspects of the vomeronasal system in canids. Specifically, we focus on the Iberian wolf (*Canis lupus signatus*), an emblematic species due to its ecological importance, cultural significance, and role in ecosystem dynamics. Surprisingly, to the best of our knowledge, the neuroanatomical features of its vomeronasal system, have remained largely unexplored until now. Notably, research on VNS in domestic dogs has seen a remarkable increase in the last decade, highlighting its crucial role in shaping the sociosexual behaviors of domestic canines (Muñiz-de Miguel et al. 2023), but also its potential involvement in pathological conditions that result in significant behavioral changes (Asproni et al. 2016). As a consequence, clinical interest in this sensory system has substantially increased (Pageat and Gaultier 2003; Dzięcioł et al. 2020).

Contrastingly, one of the primary limitations in the existing literature is the paucity of research concerning wild or feral canids. Only a few notable exceptions exist such as the study by Chengetanai et al. (2020), which addressed the neuroanatomy of the African wild dog AOB within their broader investigation of the olfactory system of this species, or the recent contributions in foxes, have unveiled considerable anatomical and functional differences when compared to their domestic counterparts. It Is remarkable how these fox studies have found specific features in the structure and neurochemistry of the VNO (Ortiz-Leal et al. 2020), AOB (Ortiz-Leal et al. 2022b), and in the transition zone commonly referred to as the olfactory limbus (Ortiz-Leal et al. 2023).

In the subsequent sections, we will focus on the key findings of our research, specifically addressing the unique neuroanatomical features of the VNS in wolves and its evolutionary implications. To accomplish this, we will not only compare our data with the extensively researched VNS of the domestic dog, but also incorporate available data from other wild canid species. Additionally, our goal is to contextualize our results within the larger framework of carnivorous taxa, specifically with a focus on felids, mustelids, and ursids, to further elucidate the adaptive, evolutionary, or potentially convergent characteristics of these chemosensory systems. This study thereby seeks to provide a neuroanatomical basis for guiding future investigations into the chemical ecology of not only wild canids but also other carnivorous taxa.

### VNO macroscopic study

Utilizing both cross-sectional macroscopic anatomy and computed tomography (CT) scans, we were able to delineate the topographic relationships and macroscopic features of the wolf VNO with remarkable precision. The use of CT scans to characterize VNO anatomy has been relatively limited, with only a small number of studies focusing specifically on this area. Previous research has been largely restricted to goats (Moawad et al. 2017), camels (Alsafy et al. 2014), bats (Yohe et al. 2018), and mice (Mucignat 2004; Levy et al. 2020), the latter employing high-resolution magnetic resonance and micro-CT techniques. Our results corroborate that the wolf VNO is situated bilaterally in the most rostral and ventral region of the nasal cavity. It lies laterally to the vomer bone and ventrally to the cartilage of the nasal septum, mostly accommodating in the internal side of the palatine fissure. The serial-anatomical sections show how the VNO is highly adapted to the contours of the nasal cavity, reinforcing the idea that its strategic placement may optimize its functional efficacy, allowing an easy communication through the incisive duct with the external environment through both the nasal and oral cavities. This hints at a complex interplay between the nasal and oral cavities, potentially facilitating a multi-modal sensory input for the wolf.

Due to its intricate location and the fact that the cartilaginous capsule of the VNO is entirely covered by the respiratory mucosa of the nasal cavity, visualizing the organ is challenging both in vivo and post-mortem. The presence of the caudal nasal myelinated nerve at the most caudal extremity of the VNO serves as a reliable indicator for its location. Additionally, the vomeronasal nerve fibers were observed to run in a caudo-dorsal direction, suggesting an integrated neuroanatomical pathway with the main olfactory system. The cartilaginous capsule was found to almost entirely envelop the parenchyma of the VNO, except for its dorsolateral part. This is similar to what has been described in other carnivores such as dogs (Salazar et al. 2013), foxes (Ortiz-Leal et al. 2020), ferrets (Kelliher et al. 2001), minks (Salazar et al. 1994a), and bears (Tomiyasu et al. 2017). However, it is somewhat less extensive than what has been observed in felines, where the capsule completely encloses the organ (Salazar et al. 1995). While our macroscopic study was primarily anatomical in nature, the observations have functional implications. For instance, the location of the VNO adjacent to the root of the upper canine tooth suggests a potentially critical role in the sensing of pheromones during aggressive or mating behaviors.

### VNO histological features

Our histological analysis of the VNO reveals a complex microanatomy that is critical for understanding its potential physiological and behavioral functions. The U-shaped cartilaginous structure that encapsulates the soft tissue serves a crucial role; it prevents the soft tissue from collapsing under the negative pressure generated by the vomeronasal pumping mechanism designed to intake pheromones (Meredith and O’Connell 1979; Meredith et al. 1980; Meredith 1994). The presence of a complex venous network within the wolf VNO is a noteworthy finding. Although this feature is also characteristic of the VNO in dogs and cats, as reported by Salazar et al. (1997; 2013), all the wolf specimens we studied showed a more advanced development of the vascular component. This was particularly evident in the predominance of large, muscular veins located in the dorsal and lateral regions of the organ. This venous preponderance plays a crucial role in the functioning of the vascular pump. When these vascular structures within the soft tissue contract, the lumen of the vomeronasal duct expands, creating a vacuum effect that draws in chemical molecules. On the other hand, vascular dilation causes the duct to constrict, leading to the expulsion of its contents, as described by Eccles (1982).

The limited presence and reduced size of arteries in the VNO could indicate that the organ does not require a high supply of oxygenated blood for its primary function of chemoreception. This may suggest an energy-efficient mechanism, where the organ operates at optimal functionality without needing substantial blood flow. This could also reflect evolutionary adaptations that prioritize efficiency in sensory organs (Niven and Laughlin 2008).

Our observations of blood capillaries in the neuroepithelium of the organ suggest the existence of mechanisms that enhance the efficiency of nutrient and gas supply to the VNO. Interestingly, these capillaries are in direct contact with the neuroreceptor layer, leading to speculation about the possible existence of hematogenic olfaction (Bednar and Langfelder 1930). Although this olfactory paradigm is far from proven, recent discoveries indicating that the VNO serves as a critical sensor for hemoglobin in rodents (Osakada et al. 2022) could support this hypothesis. The intra-epithelial capillaries have been previously characterized in rats (Breipohl et al. 1981), a species with notably thick epithelium requiring substantial blood supply. In contrast, intra-epithelial blood vessels have not been reported in species where this neuroepithelium is composed of only a few cell rows as, for example, in the lemur (Smith et al. 2007; Smith et al. 2015), the tree shrew and the slow loris (Loo and Kanagasunteram 1972), certain primates (Smith et al. 2011a), and bats (Bhatnagar and Meisami 1998; Bhatnagar and Smith 2007). Interestingly, the presence of these intra-epithelial capillaries in wolves, a species with significantly fewer neuroreceptor cells compared to rodents, is quite striking. To the best of our knowledge, no descriptions of the dog VNO refer to the presence of such intraepithelial capillaries (Dennis et al. 2003; Salazar et al. 2013) . It could represent a significant morphological distinction that may have implications for the functional capabilities VNO in these closely related species.

The interaction of the vomeronasal duct with the external environment shows notable similarities between what we have observed in our decalcified histological series in wolves and what is described in dogs (Adams and Wiekamp 1984; Salazar et al. 2013). This indirect interaction involving the incisive duct is also observed in other carnivores such as foxes (Ortiz-Leal et al. 2020), minks (Salazar et al. 1994a), and bears (Tomiyasu et al. 2017), as well as in mammals from other orders including cows (Jacobs et al. 1981), moose (Vedin et al. 2010), and hedgehogs (Kondoh et al. 2021). One end of the incisive duct communicates with the vomeronasal duct through the ventral recess of the nasal cavity, while its other end connects to the oral cavity via the incisive papilla. This distinct anatomical configuration differentiates them from rodents and lagomorphs. In the latter groups, the vomeronasal duct directly opens into the nasal cavity, and the incisive duct serves as an independent link between both cavities (Vaccarezza et al. 1981; Villamayor et al. 2018).

The relatively sparse glandular tissue, concentrated near the ventral and particularly the dorsal commissures, appears to play a specialized role in either secretion or absorption, thus maintaining a continuous mucous environment within the vomeronasal duct (Halpern and Martínez-Marcos 2003). In wolves, as is the case in dogs (Kondoh et al. 2020) and foxes (Ortiz-Leal et al. 2020), few glands are observed in the central and medial portions of the organ; however, they become progressively more numerous in the caudal portions. Using PAS and Alcian Blue stains, we were able to characterize the nature of the glandular secretions in the wolf VNO as both PAS- and AB-positive. This dual nature of PAS- and AB-positive vomeronasal glands has also been observed in foxes (Ortiz-Leal et al. 2020). In contrast, Kondoh et al. (2020) reported that vomeronasal glands in dogs were solely PAS-positive. Regarding other Carnivore species, Tomiyasu et al. (2018) found both PAS- and AB-positive glands in bears, while in cats and dogs, only PAS-positive glands were identified by Salazar et al. (1996) and Kondoh et al. (2020), respectively. These differences may be attributed to the specific region of the VNO examined, as most studies have focused only on its central region where we have found the glandular tissue density to be much lower. Future studies that examine the nature of vomeronasal gland secretion along the entire axis of the dog VNO should help to clarify this issue.

Other histological features of the wolf VNO are comparable to the information available for dogs. Specifically, the abundant connective tissue found throughout the wolf VNO likely plays a critical role in maintaining structural integrity. This tissue acts as a scaffold, helping to preserve the intricate microanatomy that is essential for the specialized functions of the VNO (Takami 2002). Additionally, the identification of two distinct types of nerve fibers in the VNO— myelinated and unmyelinated—is common to both species. These are respectively related to sensory perception and autonomic control of blood vessels and glands (Iwanaga and Nio-Kobayashi 2020).

### VNO immunohistochemical features

The immunohistochemical characterization of the wolf VNO reveals distinct patterns of expression for different types of G-protein subunits and other neural markers, such as calbindin, calretinin, OMP, and GAP-43. These markers serve as indicators for various functional roles and developmental stages of the neuroreceptor cells within the VNO.

Immunohistochemical analysis employing specific antibodies against the alpha-subunits of Gi2 and Go proteins holds particular significance. This assertion is backed by both neurochemical (Shinohara et al. 1992) and genomic studies (Dulac and Axel 1995; Herrada and Dulac 1997; Matsunami and Buck 1997; Ryba and Tirindelli 1997) in rodents, which have consistently indicated that the Gαi2 protein is tied to the expression of the V1Rs receptor family in the VNS, whereas the Gαo protein is linked to the V2Rs family. Subsequent research revealed the absence of the Gαo pathway in various mammals, including both Laurasiatheria and Primates (Takigami et al. 2000; Suárez et al. 2011a). However, the sparse studies focusing on G protein expression in the VNS of Carnivora have been a point of debate. Firstly, Dennis et al. (Dennis et al. 2003) reported immunopositive labelling in the dog VNO neurosensory epithelium using both anti-Gαi2 and anti-Gαo antibodies. They speculated that this unexpected result could be an unintended consequence of the antigen retrieval process. This theory gained further credence in a later study by Salazar et al. (2013), who reported immunonegative labelling using the anti-Gαo antibody when antigen retrieval was not applied. However, more recently, the presence of Gαo protein immunoreactivity was confirmed in the VNO neuroepithelium (Ortiz-Leal et al. 2020) and the vomeronasal nerves in the nasal mucosa and the cribriform plate (Ortiz-Leal et al. 2022b). This unexpected expression pattern in the fox has now also been confirmed in another wild canid, the wolf. Specifically, the anti-Gαo antibody mainly labelled neurons located in the basal layers of the vomeronasal neuroepithelium. These marked cells were found in proximity to intraepithelial capillaries, suggesting they may play a role in vascular interactions or hematogenic olfaction. This stands in contrast to cells labelled with the anti-Gαi2 antibody, which were situated in the epithelium central zone and showed no relation to intraepithelial capillaries.

While the immunohistochemical identification of Gαo is often hailed as a reliable marker for V2R expression in the VNO, this notion is not fully corroborated by existing genomic studies. These studies suggest that functional V2R genes have become vestigial in numerous mammalian groups, including Carnivores, through a process of accelerated pseudogenization (Young and Trask 2007). However, translating genomic findings into neuroanatomical facts presents challenges. There is a growing body of work highlighting the mismatch between genetic and morphological aspects in chemosensory systems. To bridge this gap, further morphological investigation is essential, particularly focusing on the associated brain regions, glands, and ducts (Yohe and Krell 2023). The high incidence of pseudogenes among vomeronasal receptors raises questions and could account for the disparity between sequencing and anatomical findings. Evidence of this complexity is found in an olfactory receptor gene that, despite having a premature stop codon, encodes a functional protein due to effective translational read-through (Prieto-Godino et al. 2016; Stensmyr 2016). Additionally, transcriptomic studies have detected the expression of vomeronasal pseudogenes in the mouse VNO (Oboti et al. 2015; Dietschi et al. 2022).

The possibility that the Gαo protein may serve a function in cell-to-cell signaling within the wolf neuroepithelium cannot be dispeleld, even though such a role would be without precedent in the mammalian vomeronasal neuroepithelium. This hypothesis does not align with the immunolabelling patterns seen in the wolf VNO, which extend through dendritic processes, cell bodies, and axons forming the vomeronasal nerves—a pattern that is consistent with both G proteins being implicated in transduction mechanisms (Mohrhardt et al. 2018).

The detection of both Gαi2 and Gαo proteins in the sensory epithelia of the wolf and fox VNO contrasts with the isolated expression of Gαi2 protein in other Carnivores like dogs or cats (Salazar and Sánchez-Quinteiro 2011; Salazar et al. 2013). This raises intriguing questions about the impact of domestication. The lack of Gαo protein expression in the VNS of domestic animals like goats (Takigami et al. 2000), sheep (Salazar et al. 2007), dogs (Salazar et al. 2013), and cats (Salazar and Sánchez-Quinteiro 2011) has led to the hypothesis that domestication may have led to the degeneration of the VNS (Jezierski et al. 2016).

A range of supplementary antibodies were employed for the immunohistochemical study of the VNO, including anti-CB, anti-CR, anti-GAP-43, and anti-OMP. The anti-CB and anti-CR antibodies has been frequently used to characterize neuronal subpopulations, revealing unique expression profiles in the VNS across different species (Bastianelli and Pochet 1995; Malz et al. 2000; Briñón et al. 2001). In wolves, the anti-CB antibody demonstrated a distinct immunolabelling pattern in the sensory epithelium, highlighting a subset of neuroreceptor cells predominantly located in the deeper layers of the epithelium. The labelling was most concentrated in the cell bodies and less so in the dendritic extensions. Conversely, the anti-CR antibody produced labelling that complemented that of the anti-CB antibody, targeting somata more superficially located within the epithelial layer. Notably, dendritic buttons on these neurons appeared bulb-shaped and were strongly stained.

The anti-GAP43 antibody, employed to identify neurons undergoing axonal development and synaptogenesis (Verhaagen et al. 1989; Gispen et al. 1991; Ramakers et al. 1992), showed intense and widespread labelling. This pattern was consistent with findings in both fox (Ortiz-Leal et al. 2020) and dogs (Dennis et al. 2003), suggesting that active neuronal regeneration is occurring in the canine vomeronasal sensory epithelium. This ongoing plasticity could be in response to the VNO being frequently exposed to various environmental substances that have the potential to damage cellular structures (Ogura et al. 2010). These observations underscore the significance of the vomeronasal sensory system for canids.

Finally, the anti-OMP antibody targeted the olfactory marker protein, which is expressed in mature neurons in both the main olfactory system and the VNS (Farbman and Margolis 1980). It has shown immunopositive labelling in a range of species, including rats (Rodewald et al. 2016), mice (Mechin et al. 2021), mole rats (Dennis et al. 2020), and primates (Smith et al. 2011b). The ubiquitous presence of OMP in the wolf VNO suggests a uniform stage of neuronal maturation throughout the epithelium.

Beyond the elaborate neurochemical pattern observed in the sensory neuroepithelium of the wolf VNO—a reflection of the complex interplay of molecular, cellular communication, and signal transduction mechanisms at work in pheromonal information processing—the immunohistochemical study has allowed us to identify a striking, distinctive feature of the wolf VNO. This feature consists of abundant clusters of neuronal cells situated in the most basal part of the sensory neuroepithelium. These clusters extend several hundreds of micrometers immediately beneath the basal layer of neuroreceptor cells. They are composed of oval-shaped cells with large, spherical nuclei, closely packed together and seemingly devoid of processes. While their morphology superficially resembles that of the neuroreceptor cells of the neuroepithelium, these cells display a specific neurochemical pattern. On one hand, they are OMP-positive, reinforcing their role in olfactory signal transduction. On the other hand, they are immunonegative for GAP-43, suggesting that they exist in a state of lower differentiation compared to the highly GAP-43-positive neuroreceptor cells. This observation does not appear to support the notion that these are undifferentiated cells serving to renew the epithelium, akin to typical basal cells. Regarding calcium-binding proteins, these clusters are calretinin-positive and calbindin-negative, which could initially establish a link between them and the calretinin-positive cell subpopulation.

The presence of these clusters seems to be specific to the wolfs VNO. Our comprehensive study of the fox VNO, using similar markers and protocols, did not reveal this type of neuroepithelial organization, nor did studies in other carnivores such as dogs, minks, ferrets, or bears. Even across decades of studying the VNO in a diverse array of mammals, we have not encountered a similar neuroepithelial organization (Torres et al. 2023b). Therefore, more specific research on these cell clusters, employing additional markers or even genomic approaches such as single-cell technology, are necessary to further characterize this unique cell population, thereby enhancing our understanding of its functional role or significance.

### VNO lectin histochemical labelling

Our lectin histochemical investigation reveals subtle differences in the labelling patterns of UEA and LEA lectins within the VNO sensory epithelium and vomeronasal nerves. Both lectins produce positive labelling in neuroreceptor cells and the vomeronasal nerves, though they diverge significantly when it comes to the basal neuroepithelial cell clusters. LEA labels these neuronal clusters, while UEA does not, suggesting distinct molecular interactions between these lectins and the cellular components of the clusters. Furthermore, LEA shows stronger labelling in the apical processes, while UEA is more concentrated in basal areas of the epithelium, although it never labels the basal clusters.

In the respiratory epithelium, both lectins exhibit diffuse but specific labelling patterns. LEA primarily targets the apical processes, whereas UEA labelling is sparser, with a few strongly labelled cells scattered throughout the epithelium. Finally, within the mucociliary complex, UEA shows stronger labelling compared to LEA. The distinct labelling patterns imply that the glycoconjugates recognized by these lectins could have specialized molecular functions within the VNO.

### AOB macroscopic and microscopic anatomy

To the best of our knowledge, this study represents the first morphological investigation of the wolf AOB. As is the case in other canids studied, such as the dog (Nakajima et al. 1998; Salazar and Sánchez-Quinteiro 2011), the African wild dog (Chengetanai et al. 2020) and the fox (Ortiz-Leal et al. 2022b), the AOB is very small in size compared to the MOB, making its macroscopic identification quite challenging. Only through the use of serial histological sections can this structure be accurately localized. The study of the cytoarchitecture of the wolf AOB is particularly important, given that there has been ongoing debate for decades about the moderate development, and even the very existence, of the AOB in dogs. This controversy was only resolved in the 1990s with the advent of lectin histochemical staining techniques (Salazar et al. 1992; Salazar et al. 1994b).

Our research in the wolf corroborates the existence of an AOB with dimensions comparable to those found in domestic dogs. However, the wolf AOB exhibits a more pronounced laminar organization. Notably, there is a significant development of the superficial nervous and glomerular layers of the AOB. Notably, the wolf AOB features a higher number of mitral cells, a principal cell type rarely evident in histological sections of the dog AOB. As a result, the wolf AOB can be described as having a well-defined mitral-plexiform layer. Our findings are in line with the comprehensive study by Chengetanai et al. (2020) in the African wild dog olfactory system which included an examination of the AOB. While the size and development of the AOB in wild canids may appear to be limited, this is not the case when compared to other groups of carnivores. Specifically, mustelids such as the mink (Salazar et al. 1998) and ferret (Kelliher et al. 2001), as well as herpestids like the meerkat (Torres et al. 2021), possess poorly differentiated AOBs. Altogether, this morphological research corroborates the existence of distinct lamination patterns in the AOB across wild canid populations and supports the theory that the selection pressure linked to domestication may have had a regressive impact on the degree of differentiation in the dog vomeronasal system, a sensory pathway crucial for survival in the wild.

### AOB immunohistochemistry and lectin histochemistry

The neurochemical pattern of the wolf AOB in its superficial, nervous, and glomerular layers is consistent with what is observed in the VNO. That is, the proteins Gαi2, CB, CR, and OMP, expressed in the vomeronasal neuroepithelium and in the vomeronasal nerves both in the VNO and the nasal mucosa, are also reaching the superficial layers of the AOB where they produce an immunopositive labelling. However, the superficial layers are Gαo negative. This implies that the Gαo positive vomeronasal neuroreceptors are projecting their information to different areas of the olfactory bulb. This pattern is identical to the one observed in the VNS of the fox (Ortiz-Leal et al. 2022b), a species in which a recent study suggest that the projection of the Gαo vomeronasal afferents takes place to the transition zone known as the olfactory limbus (Ortiz-Leal et al. 2023). The investigation of the possible presence of a similar olfactory limbus in the wolf is beyond the objectives of this work, but it is a matter worthy of further study. Of all the markers described, Chengetanai et al. (2020) in their study of the African wild dog AOB only employed anti-CR, obtaining a pattern similar to the one described by us in the wolf. Furthermore, we have used in our study of the wolf AOB antibodies against MAP-2 and GFAP proteins, which have allowed us to characterize the remarkable development of the dendritic tree of the mitral-plexiform layer and the glial component both at the level of the enveloping glia and astrocytes.

Both UEA lectin and LEA produce immunopositivity in the superficial areas of the AOB, but while the former is specific to the AOB, the latter stains both the AOB and the MOB. This pattern is identical to the one observed in the dog (Salazar et al. 1992; Salazar et al. 2013) and the fox (Ortiz-Leal et al. 2022b), and it confirms the usefulness of UEA as a marker for the VNS of canids and LEA as a general marker for the VNS and MOS in the same family. It is significant, however, that the BSI-B_4_ lectin, specific to the VNS of the rat (Ichikawa et al. 1992), is nevertheless negative in the case of the wolf, as it is a demonstration of the extremely high specificity that the expression of glycoconjugates has in different groups of mammals.

In conclusion, this comprehensive study of the wolf VNS (vomeronasal organ, vomeronasal nerves, and accessory olfactory bulb) has provided the first detailed characterization of its macroscopic anatomy, histology, and neurochemical and histochemical profile. Our findings highlight significant differences between the wolf *Canus lupus signatus* and its domestic counterpart, *Canis lupus familiaris*, in both structural and neurochemical terms. This supports the hypothesis that the domestication of the dog ancestor has led to a regression of specific molecular and neurochemical features within the vomeronasal system. Beyond the evolutionary implications, it is noteworthy that the wolf VNS exhibits unique characteristics. It aligns with other animal models showing dual expression of G proteins subunits in their VNO, a finding of great interest. Particularly noteworthy is the presence of extensive neural clusters in the VNO neuroepithelium, previously undocumented in other canid species. This adds new depth to our comparative understanding of the mammalian vomeronasal system. Future molecular and genomic investigations should shed light on the meaning of this structural and neurochemical features.

**Table 1.**
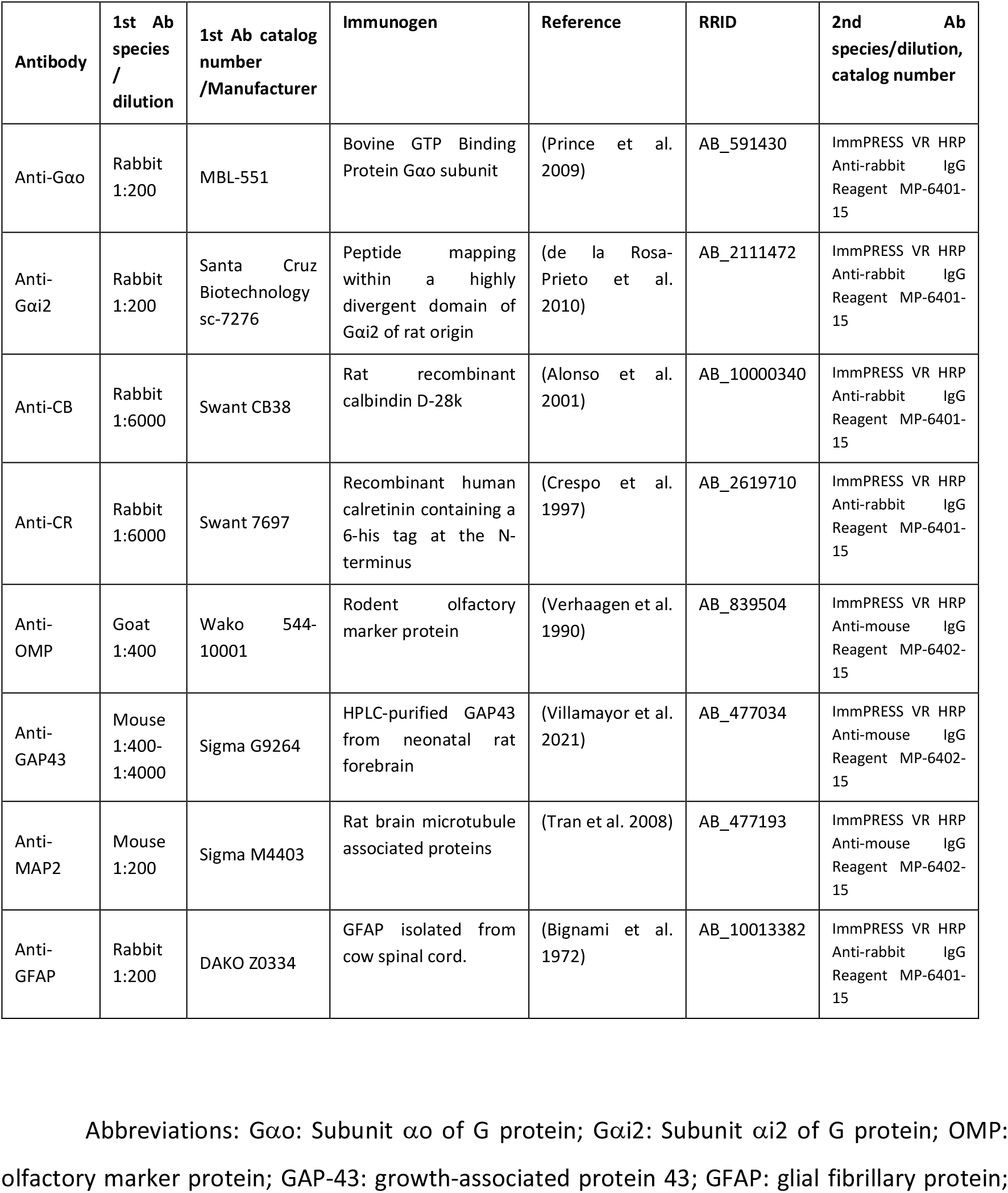
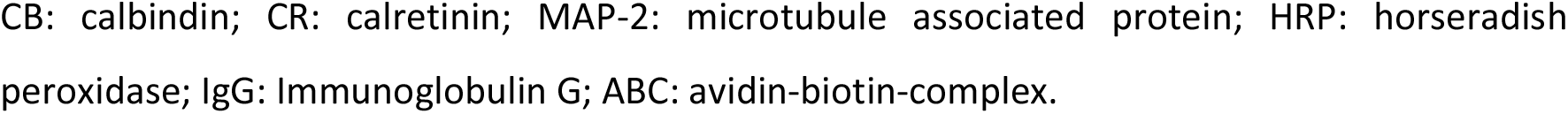
Detailed information on the antibodies used in this study: Species of elaboration, dilution, catalogue number, manufacturer, target immunogens, relevant reference for each antibody, RRID codes, and secondary antibody employed. Specifications for each antibody used in the study, encompassing supplier information, dilution ratios, target immunogens, and Research Resource Identifiers (RRID). In all cases, the immunoreactivity patterns observed in wolf samples were consistent with those previously documented across various mammalian species. Relevant references for each antibody are included.

| **Table 1. Detailed information on the antibodies used in this study: Species of elaboration, dilution, catalogue number, manufacturer, target immunogens, relevant reference for each antibody, RRID codes, and secondary antibody employed.** |  |  |  |  |  |  |
| --- | --- | --- | --- | --- | --- | --- |
| **Antibody** | **1st Ab species / dilution** | **1st Ab catalog number /Manufacturer** | **Immunogen** | **Reference** | **RRID** | **2nd Ab species/dilution, catalog number** |
| Anti-Gαo | Rabbit 1:200 | MBL-551 | Bovine GTP Binding Protein Gαo subunit | (Prince et al. 2009) | AB_591430 | ImmPRESS VR HRP Anti-rabbit IgG Reagent MP-6401-15 |
| Anti-Gαi2 | Rabbit 1:200 | Santa Cruz Biotechnology sc-7276 | Peptide mapping within a highly divergent domain of Gαi2 of rat origin | (de la Rosa-Prieto et al. 2010) | AB_2111472 | ImmPRESS VR HRP Anti-rabbit IgG Reagent MP-6401-15 |
| Anti-CB | Rabbit 1:6000 | Swant CB38 | Rat recombinant calbindin D-28k | (Alonso et al. 2001) | AB_10000340 | ImmPRESS VR HRP Anti-rabbit IgG Reagent MP-6401-15 |
| Anti-CR | Rabbit 1:6000 | Swant 7697 | Recombinant human calretinin containing a 6-his tag at the N-terminus | (Crespo et al. 1997) | AB_2619710 | ImmPRESS VR HRP Anti-rabbit IgG Reagent MP-6401-15 |
| Anti-OMP | Goat 1:400 | Wako 544-10001 | Rodent olfactory marker protein | (Verhaagen et al. 1990) | AB_839504 | ImmPRESS VR HRP Anti-mouse IgG Reagent MP-6402-15 |
| Anti-GAP43 | Mouse 1:400-1:4000 | Sigma G9264 | HPLC-purified GAP43 from neonatal rat forebrain | (Villamayor et al. 2021) | AB_477034 | ImmPRESS VR HRP Anti-mouse IgG Reagent MP-6402-15 |
| Anti-MAP2 | Mouse 1:200 | Sigma M4403 | Rat brain microtubule associated proteins | (Tran et al. 2008) | AB_477193 | ImmPRESS VR HRP Anti-mouse IgG Reagent MP-6402-15 |
| Anti-GFAP | Rabbit 1:200 | DAKO Z0334 | GFAP isolated from cow spinal cord. | (Bignami et al. 1972) | AB_10013382 | ImmPRESS VR HRP Anti-rabbit IgG Reagent MP-6401-15 |

Abbreviations: Gαo: Subunit αo of G protein; Gαi2: Subunit αi2 of G protein; OMP: olfactory marker protein; GAP-43: growth-associated protein 43; GFAP: glial fibrillary protein;
CB: calbindin; CR: calretinin; MAP-2: microtubule associated protein; HRP: horseradish peroxidase; IgG: Immunoglobulin G; ABC: avidin-biotin-complex.

## AUTHOR CONTRIBUTIONS

I.O.L., M.V.T., and P.S.Q designed the research and wrote the paper. P.S.Q., I.O.L., J.D.B.V., M.V.T., L.F., and A.L.B performed the work, analysed and discussed the results.

## Notes

### Competing Interest Statement

The authors have declared no competing interest.

